# Transferable Coarse Graining via Contrastive Learning of Graph Neural Networks

**DOI:** 10.1101/2023.09.08.556923

**Authors:** Justin Airas, Xinqiang Ding, Bin Zhang

## Abstract

Coarse-grained (CG) force fields are essential for molecular dynamics simulations of biomolecules, striking a balance between computational efficiency and biological realism. These simulations employ simplified models grouping atoms into interaction sites, enabling the study of complex biomolecular systems over biologically relevant timescales. Efforts are underway to develop accurate and transferable CG force fields, guided by a bottom-up approach that matches the CG energy function with the potential of mean force (PMF) defined by the finer system. However, practical challenges arise due to many-body effects, lack of analytical expressions for the PMF, and limitations in parameterizing CG force fields. To address these challenges, a machine learning-based approach is proposed, utilizing graph neural networks (GNNs) to represent CG force fields and potential contrasting for parameterization from atomistic simulation data. We demonstrate the effectiveness of the approach by deriving a transferable GNN implicit solvent model using 600,000 atomistic configurations of six proteins obtained from explicit solvent simulations. The GNN model provides solvation free energy estimations much more accurately than state-of-the-art implicit solvent models, reproducing configurational distributions of explicit solvent simulations. We also demonstrate the reasonable transferability of the GNN model outside the training data. Our study offers valuable insights for building accurate coarse-grained models bottom-up.

## Introduction

Coarse-grained (CG) force fields play a pivotal role in advancing molecular dynamics (MD) simulations of biomolecules by bridging the gap between computational efficiency and biological realism.^1–35^ In these simulations, biomolecules are represented with simplified models that group several atoms into a single interaction site, reducing the computational burden without sacrificing essential structural and dynamical information. This approach is crucial for studying large and complex biomolecular systems over biologically relevant timescales, which would be infeasible with atomistic simulations. CG force fields enable researchers to explore phenomena such as protein folding, ^1–5^ protein aggregation,^10–17,36–38^ and large-scale conformational changes,^39,40^ providing insights into biological processes that are otherwise inaccessible. Understandably, there is great interest in developing methodologies for building accurate and transferable CG force fields. ^7,18–26^

Tremendous advancement in the theoretical framework provides guiding principles for developing accurate CG force fields from atomistic simulation data, i.e., the bottom-up approach. ^18–23^ For example, one can consider the mapping **M** that transforms the atomic Cartesian coordinates **r** to the CG configuration **R**. The CG energy function *W* (**R**) can be explicitly defined as

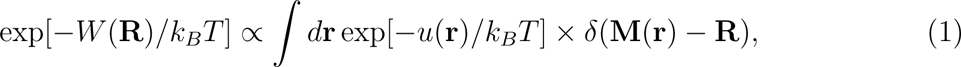

where *u*(**r**) is the potential of the assumed atomistic model. A CG force field that reproduces the potential of mean force (PMF), *W* (**R**), is referred to as *thermodynamically consistent* with the atomistic force field. ^18–23^ Being thermodynamically consistent indicates that statistical properties predicted by the CG force field will match with atomistic results.

However, despite the elegant theoretical framework, developing thermodynamically consistent CG force fields from the bottom up faces several challenges in practice. First, while pairwise potentials have found great success in explicit solvent atomistic models, ^41–43^ it has long been known that many-body effects become prominent when the degrees of freedom are reduced.^1,34,35,44–46^ The PMF is inherently a many-body potential. Owing to the lack of analytical expressions, how to best approximate *W* (**R**) to achieve the desired accuracy is unclear. Secondly, despite significant methodological developments,^22–25,44,47,48^ parameterizing CG force fields with atomistic simulation data remains nontrivial. Existing algorithms often face challenges in achieving thermodynamic consistency due to error accumulation. Often, they can only leverage small datasets due to computational cost, limiting the transferability of the resulting force field.

We propose a machine learning (ML) approach to tackle both challenges. First, we employ graph neural networks (GNNs) to represent the CG force field. GNNs trained as neural network (NN) potentials use the Cartesian coordinates of the entire molecule to calculate energy. Crucially, their high expressibility enables them to parameterize complex functional relationships, capturing many-body effects. Second, we utilize a recently developed force field optimization method, potential contrasting,^25^ to parameterize GNNs from atomistic simulation data. Potential contrasting optimizes the overlap between CG and atomistic configurational distributions, aiming to ensure thermodynamic consistency. Moreover, it is highly parallelizable and computationally efficient, allowing the use of large training sets that are essential for deriving transferable force fields.

We use the ML approach to develop transferable implicit solvent models based on 600,000 conformations from explicit solvent atomistic simulations of six proteins. An implicit solvent model is a high-resolution CG model that approximates the solvation free energy, i.e., *W* (**R**), obtained by integrating the solvent degrees of freedom.^49–51^ We demonstrate that the SchNet architecture^52^ provides a precise representation of the solvation free energy. When parameterized, the SchNet implicit solvent model, along with the gas-phase force field, can accurately reproduce the free energy profiles from explicit solvent atomistic simulations. The SchNet model performs well for proteins outside the training set, indicating its transferability. We also introduce several methodological advancements necessary for parameterizing and simulating the GNN-based force field, such as a pre-training procedure that enhances the robustness and stability of MD simulations and a re-weighting scheme that enables efficient free energy calculations. Our results will be of wide interest to researchers building accurate coarse-grained models. The methodological insights should also help advance the training of GNNs as transferable force fields.

## Results

### GNNs enable transferable learning of solvation free energy

ML techniques have proven powerful for force field learning as they capture many-body effects.^53–55^ While early ML force fields relied on hand-picked descriptors, ^56,57^ GNNs with only Cartesian coordinates and atom types as inputs have become increasingly popular for force field parameterization.^58–73^ Molecules in 3D space can be conveniently abstracted into a graph, where nodes are atoms. Edges represent covalent and noncovalent interactions^74^ (Figures 1A-C). Such a graph is naturally invariant to atom indexing, and rotational and translational symmetry of energy functions can be encoded straightforwardly.

**Figure 1:**
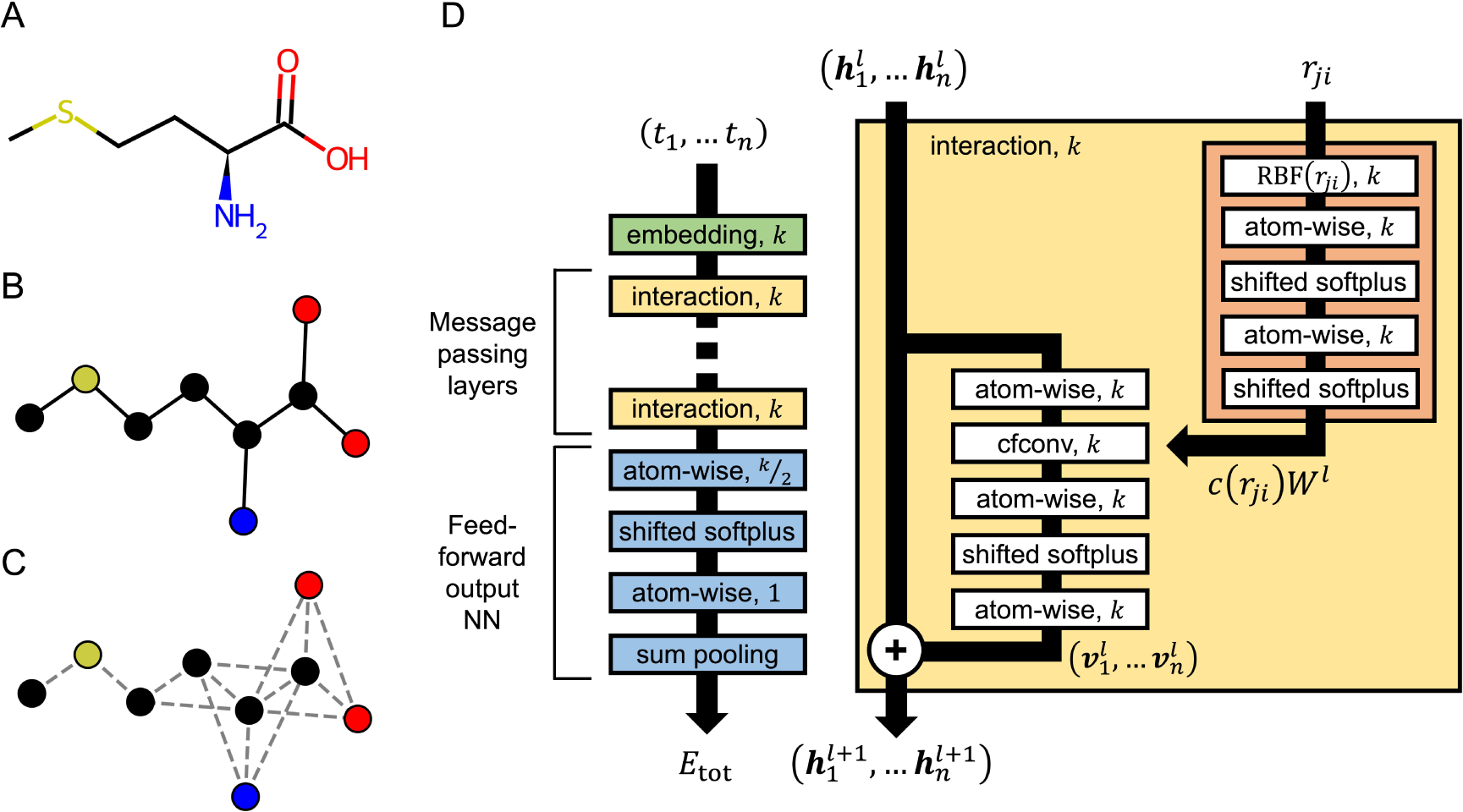
Graph neural networks allow convenient encoding of structural and chemical information of molecules for energy prediction. (A-C) Illustration of the mapping from a chemical representation of the methionine residue (A) to a graph with edges representing covalent bonds (B) and a fully connected graph of methionine with edges representing all atoms within *r*_cut_ = 2.5 Å (C). Hydrogens are omitted from the graphs for clarity. (D) Illustration of the SchNet architecture.^52^ The embedding layer (green) assigns *k* learnable parameters, 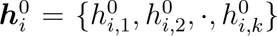, for each atom *i* in the protein based on its atom type, *t_i_*. The featurizations are updated by a series of message passing layers, i.e., interaction blocks, (yellow) that account for the chemical environment surrounding individual atoms. See the *Methods* Section for additional discussion of interaction blocks. After *N*_IB_ interaction blocks, the final featurizations, 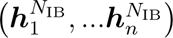, are provided to the energy-predicting NN (blue), which computes the total potential energy for the protein as 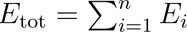.

We adopt the SchNet architecture introduced by Schütt et al. ^52^ to represent the PMF of CG models. As depicted in Figure 1D, the network architecture can be broadly divided into three parts: an embedding layer that assigns the initial featurization to each atom, multiple message-passing layers that update the featurization, and finally, a feed-forward NN that predicts an atomic contribution to the total system energy from the updated featurization. Featurization defines the characteristics associated with a specific atom type. The message-passing step is typically performed iteratively, allowing nodes to gather information from increasingly distant neighbors, thus accounting for interactions among atoms. These layers are frequently referred to as interaction blocks. Feature updates explicitly make use of structural information inferred from Cartesian coordinates. Therefore, the SchNet model resembles a typical energy function that transforms the Cartesian coordinates of the entire molecule, along with knowledge about its chemical makeup, into a single value representing the energy. Furthermore, backward propagation can compute forces for MD simulations.

Most evaluations of SchNet and other GNNs focus on organic molecules with less than 21 atoms.^52,75^ Therefore, we need to assess SchNet’s ability to predict interaction energies for much larger protein molecules. We selected six proteins: chignolin CLN025 (CLN025), Trp-cage, BBA, Villin, WW domain, and NTL9. The smallest protein in this dataset has 166 atoms. For each protein, we collected 200,000 conformations from MD simulations (see Supporting Information and Tables S1 - S3 for details) and computed the solvation free energy, *E*_GBn2_, using the GB-neck2 (GBn2) implicit solvent model.^76^ Since *E*_GBn2_ is a many-body potential, its prediction is a challenging problem for SchNet.

We explored several SchNet architectures by focusing on adjusting two hyperparameters that directly impact the message passing layers: the cutoff distance *r*_cut_ and the number of interaction blocks *N*_IB_. Here, *r*_cut_ defines the nearest neighbors connected with edges to a given atom and used to compute feature updates in each interaction block, while *N*_IB_ determines the number of iterations for feature updates. Increasing *r*_cut_ while keeping the number of interaction blocks constant at *N*_IB_ = 3 results in a continuously decreasing training root-mean-squared error (RMSE), as illustrated in Figure 2A. Next, after selecting a well-performing smaller *r*_cut_ = 1.8 nm, we explored the effect of varying *N*_IB_. Figure 2A suggests that *N*_IB_ has a less pronounced impact on training RMSE compared to *r*_cut_. For the most accurate model (*N*_IB_ = 3, *r*_cut_ = 5 nm), SchNet indeed performs well for all proteins. Higher RMSE is observed for increasingly larger proteins, as shown in Figure 2B.

**Figure 2:**
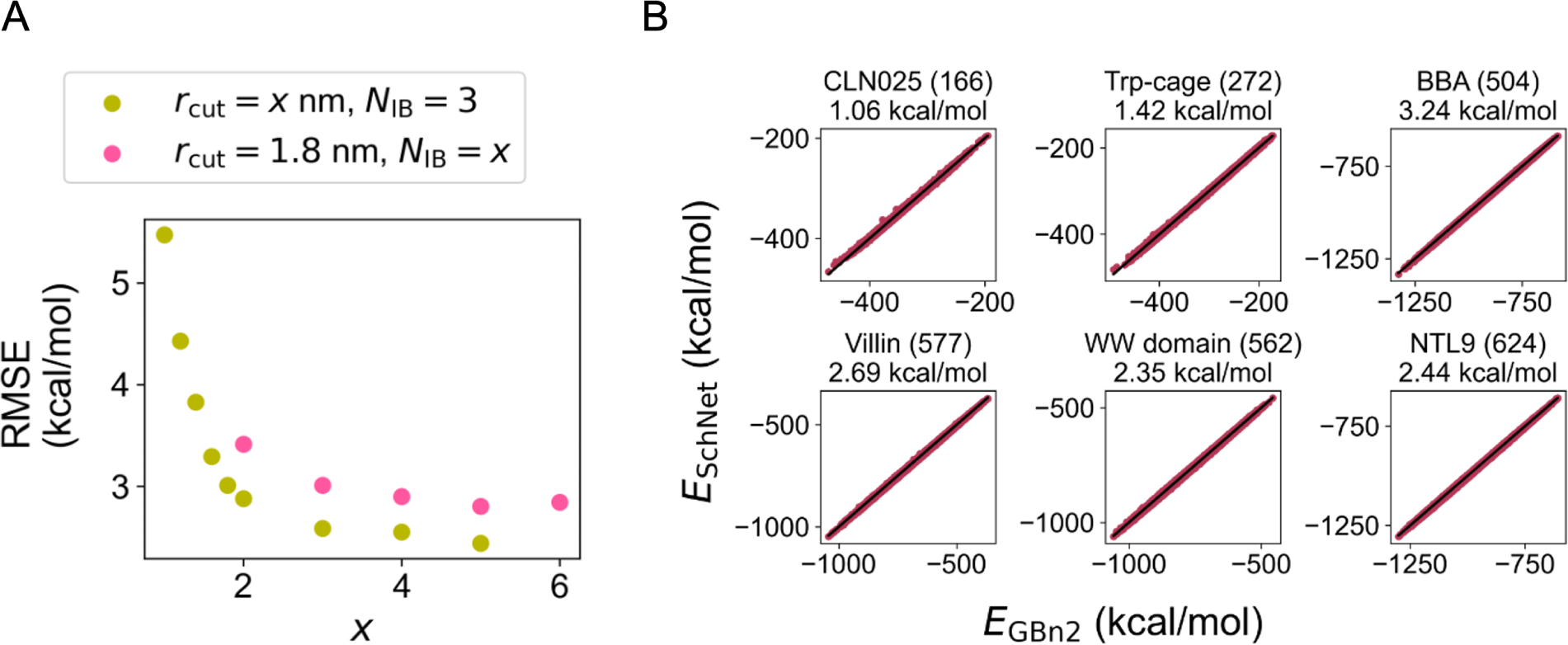
SchNet accurately reproduces the many-body solvation free energy for proteins of various sizes. (A) Average root-mean-squared error (RMSE) across all configurations and all proteins with different hyperparameters *r*_cut_ and *N*_IB_ in SchNet design. (B) Scatter plots detailing the agreement between the energy from the best performing SchNet model (*N*_IB_ = 3, *r*_cut_ = 5 nm), *E*_SchNet_, and the GBn2 solvation free energy, *E*_GBn2_, for different configurations of each protein. The number of atoms in each protein is shown in parentheses. Average RMSEs are shown for each protein.

However, as noted in previous works,^21,77^ training RMSE alone does not directly indicate the accuracy of an ML force field in simulations. Notably, the training RMSEs for the best and worst-performing hyperparameter combinations differ by only approximately 2.5 kcal/mol. To further evaluate different SchNet architectures, we devised an efficient scheme for computing another important quantity in simulations: free energy. Specifically, we computed the free energy profile as a function of the RMSD from the folded state using the SchNet model and a free energy perturbation method.^78^ Instead of conducting computationally intensive simulations with the SchNet model, we estimated the free energy using configurations sampled from RMSD-biased umbrella sampling in GBn2 implicit solvent. This scheme is detailed in the *Methods* section.

The estimated free energy profiles depicted in Figure 3 suggest that, despite a mere *∼* 2.5 kcal/mol RMSE difference, the SchNet model with *r*_cut_ = 3 nm significantly outperforms the model with *r*_cut_ = 1 nm in replicating the results from the GBn2 implicit solvent model. Conversely, despite similar training RMSEs, the SchNet model with *r*_cut_ = 5 nm and *N*_IB_ = 3 proves significantly more accurate than the model with *r*_cut_ = 1.8 nm and *N*_IB_ = 6 (Figure S1). Free energy profiles were also computed for SchNet models with *N*_IB_ ranging from 2 to 6, but little difference is observed between these models (Figure S2). These estimated profiles thus offer a more precise assessment of model performance than errors in energy alone, showcasing the utility of our re-weighting scheme for hyperparameter optimization. These results confirm that SchNet can perform quite effectively for proteins of up to 624 atoms.

**Figure 3:**
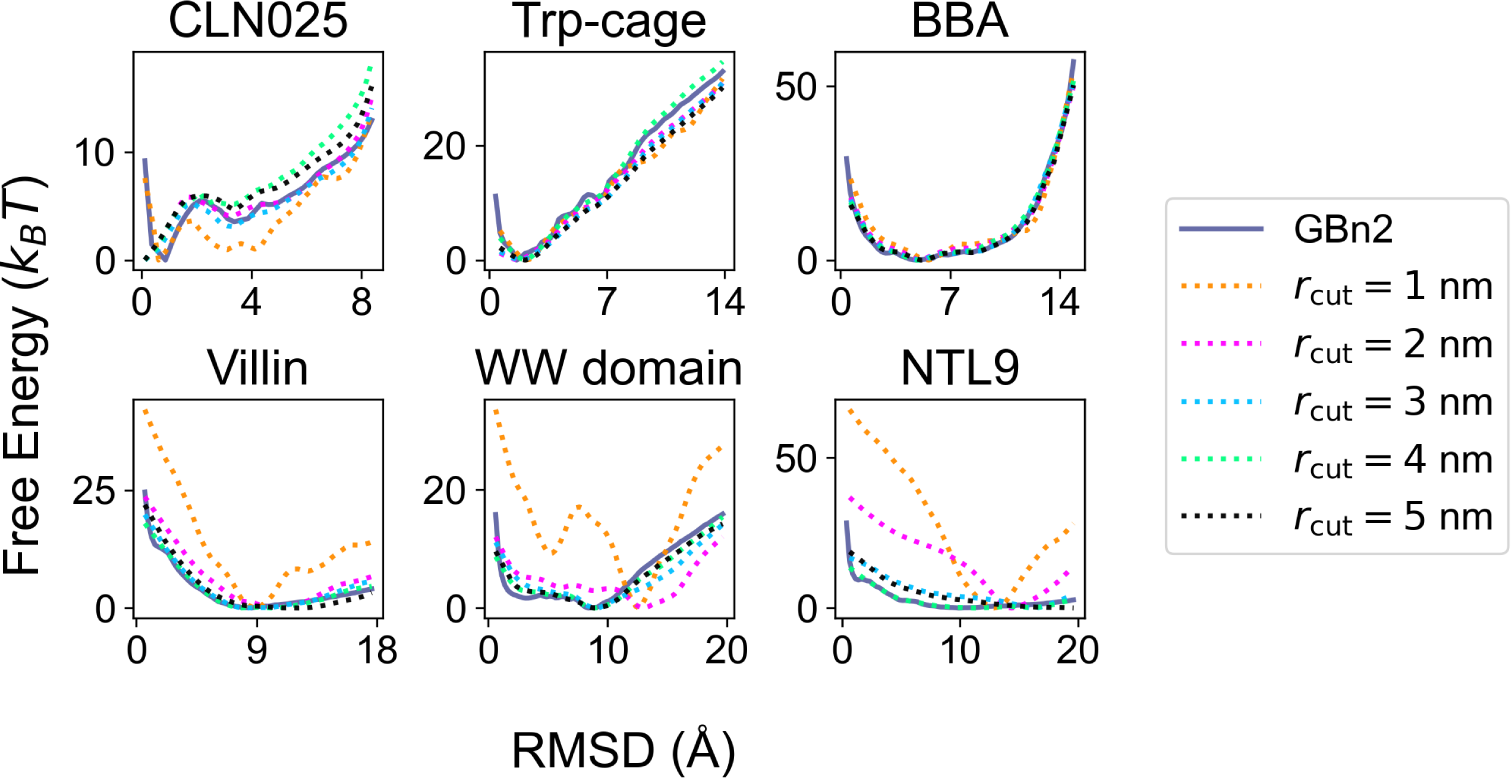
Free energy profiles for SchNet models trained to fit *E*_GBn2_ with different values of *r*_cut_. The number of interaction blocks is held constant at *N*_IB_ = 3. The *x* axes in the plots correspond to the RMSD from the folded structures. For reference, the free energy profiles computed from umbrella simulations with the implicit solvent model, GBn2, are also shown as solid lines. Results for the SchNet model were determined using a re-weighting scheme as detailed in the *Methods* Section.

### Contrastive learning of solvation free energy from explicit solvent simulations

Having demonstrated that SchNet can replicate the solvation free energy defined by the GBn2 model, our next objective is to further enhance its accuracy. Specifically, we aim to optimize SchNet to better replicate the conformational distribution obtained from explicit solvent atomistic simulations. Current implicit solvent models, including GBn2, rely on analytical expressions for the solvation free energy.^49,79–84^ While these models are computationally efficient, they often entail significant approximations, thereby limiting the achievable level of accuracy. Consequently, there is a compelling need to develop more accurate and systematically improvable representations of the solvation free energy.

We employed a method known as potential contrasting^25^ to optimize SchNet using the configurational ensemble of the six aforementioned proteins gathered from explicit solvent simulations conducted by Lindorff-Larsen et al. ^85^ (refer to Figure 4 and *Methods*). Potential contrasting, recently introduced by us, serves as a means to optimize the parameters of a CG force field by extending the noise contrastive estimation method.^86^ This approach offers computational efficiency, high parallelizability, and the ability to leverage a substantial training dataset. It’s worth noting that, due to the unavailability of the exact solvation free energy for each solute configuration, employing RMSE as a loss function is unfeasible. Consequently, the training process transitions into an unsupervised learning problem, hence our adoption of potential contrasting.

**Figure 4:**
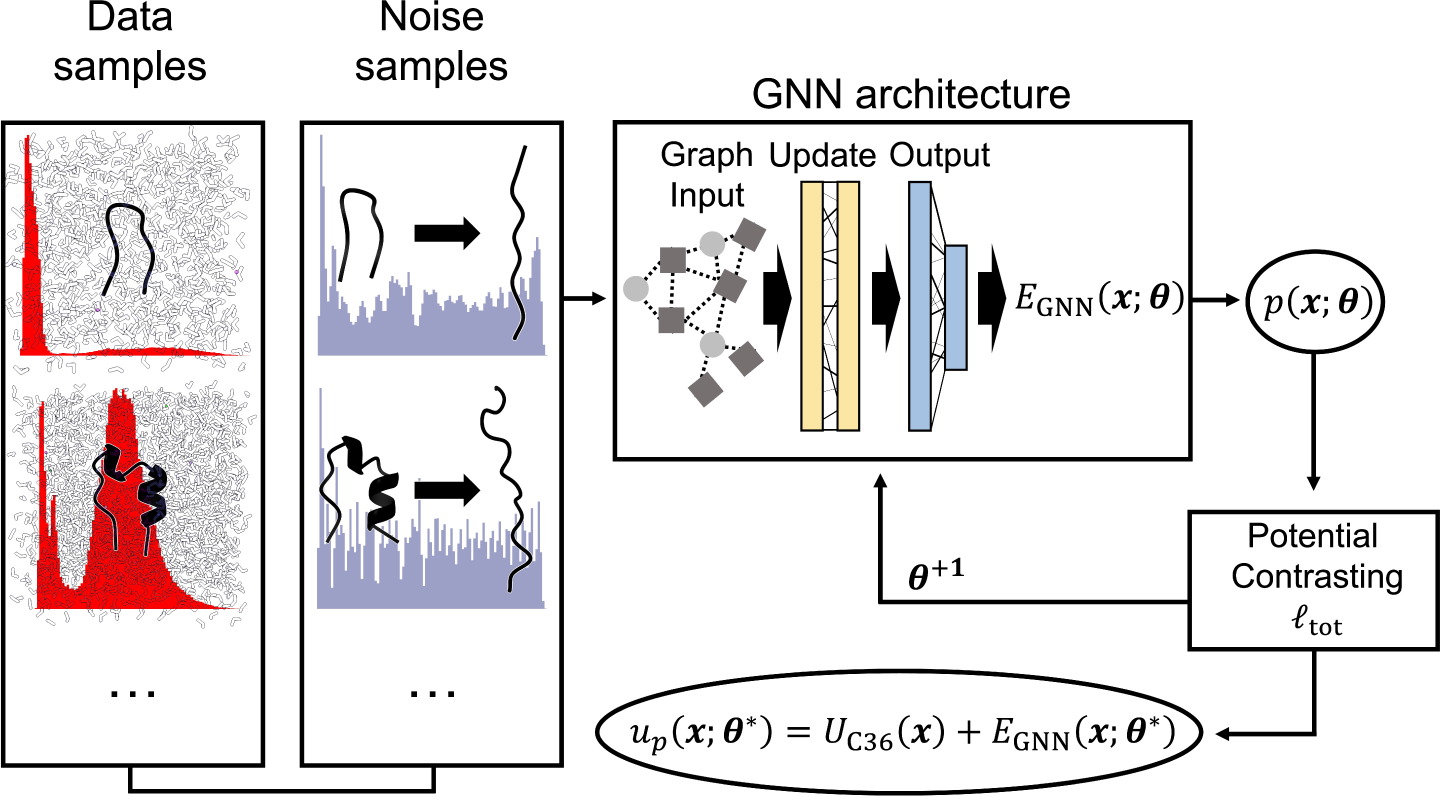
Illustration of the potential contrasting method for training SchNet to learn the solvation free energy from explicit solvent simulation data. For each protein, two ensembles of configurations are prepared to represent data and noise distributions. Data configurations are produced from unbiased long timescale explicit solvent simulations, while noise configurations are obtained from umbrella simulations using the implicit solvent model, GBn2. The Boltzmann distributions of the two ensembles are denoted as *p* (***x***; ***θ***) ∝ exp (*−β* [*u_p_* (***x***; ***θ***)]) and *q* (***x***) ∝ exp (*−β* [*u_q_* (***x***)]), respectively. *u_p_* (***x***; ***θ***) = *U*_C36_ (***x***) + *E*_SchNet_ (***x***; ***θ***), where *U*_C36_ is the gas-phase energy evaluated using the CHARMM force field^87–89^ and *E*_SchNet_ (***x***; ***θ***) is the solvation free energy, defined by the SchNet model with parameters ***θ***, for configuration ***x***. *u_q_* (***x***) represents the potential energy for the generalized ensemble constructed by combining configurations from different umbrella windows (see the Supporting Information for more details). The loss function in potential contrasting, *ℓ*_tot_, is designed to maximize *p* (***x***; ***θ***) over data samples and simultaneously minimize its values over noises. Thus, the optimal parameters ***θ^∗^*** will ensure that *p* (***x***; ***θ^∗^***) closely matches the Boltzmann distribution that the atomistic explicit solvent configurations were drawn from.

With the hyperparameters that were demonstrated to be effective for fitting SchNet to *E*_GBn2_ (*N*_IB_ = 3, *r*_cut_ = 5 nm), we observed that the SchNet model optimized using potential contrasting surpasses GBn2 by a significant margin. As illustrated in Figure 5, the free energy profiles for the SchNet model, computed with the re-weighting scheme, closely align with explicit solvent results for all six proteins.

**Figure 5:**
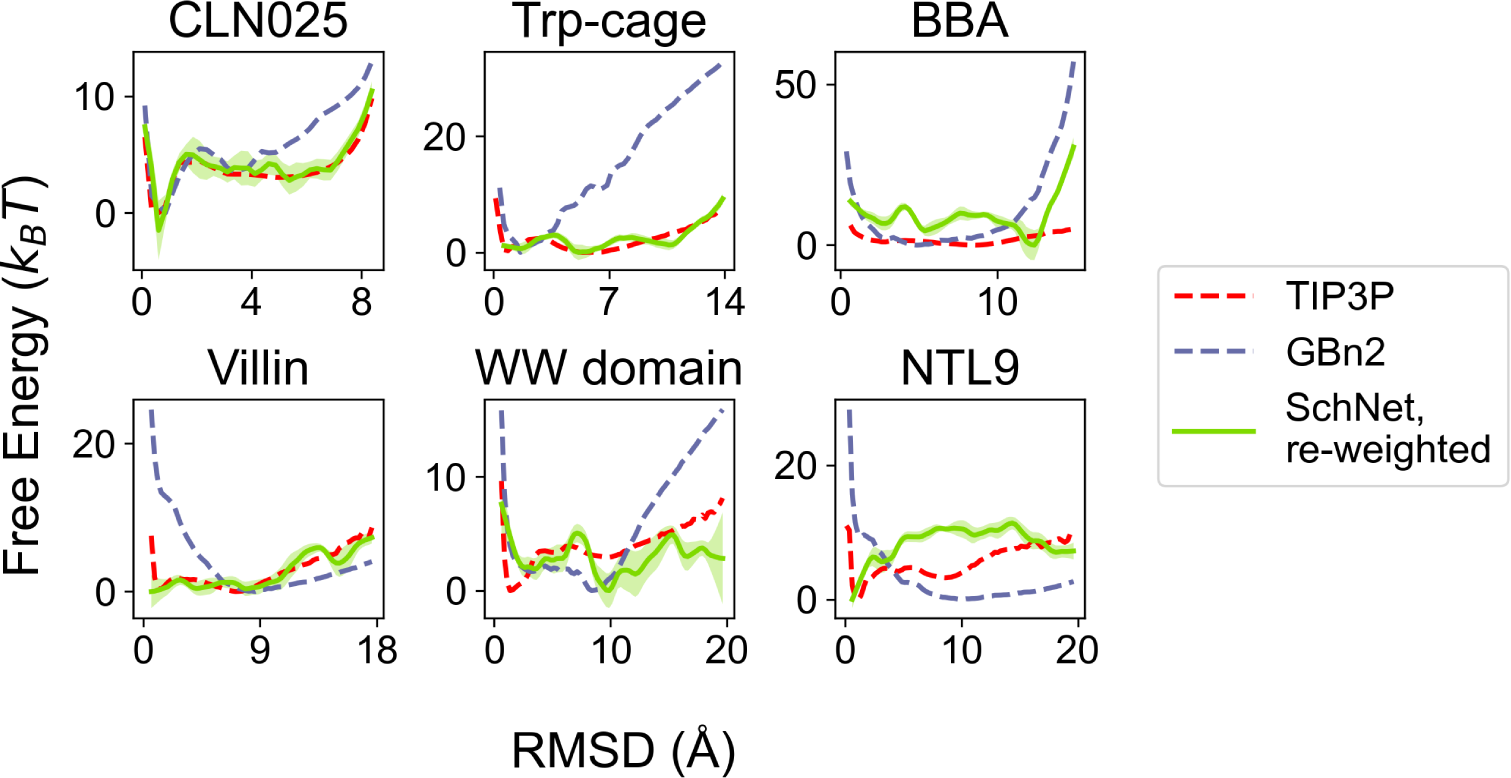
Comparison between the free energy profiles computed from explicit solvent simulations (TIP3P^90^), implicit solvent simulations (GBn2), and the optimized SchNet model with hyperparameters *N*_IB_ = 3 and *r*_cut_ = 5 nm. Due to the sharpness of the minimum in the SchNet free energy profile for CLN025, the free energy profile was uniformly shifted downwards by 1.5 *k_B_T* for better visualization. Results for the SchNet model were determined using a re-weighting scheme as detailed in the *Methods* Section.

Figure 6A illustrates that SchNet is effectively learning solvation free energies that, in comparison to GBn2, result in the destabilization of folded/compact states in favor of unfolded/extended states. It’s worth noting that this trend is reversed for the WW domain and NTL9, where GBn2 fails to capture the free energy minimum at the folded states (as shown in Figure 5). Given that it is a well-known characteristic of GB-based implicit solvents to generally over-stabilize folded/compact states, these findings support the notion that SchNet is assimilating chemically meaningful information to accurately replicate explicit solvent configurational distributions.

**Figure 6:**
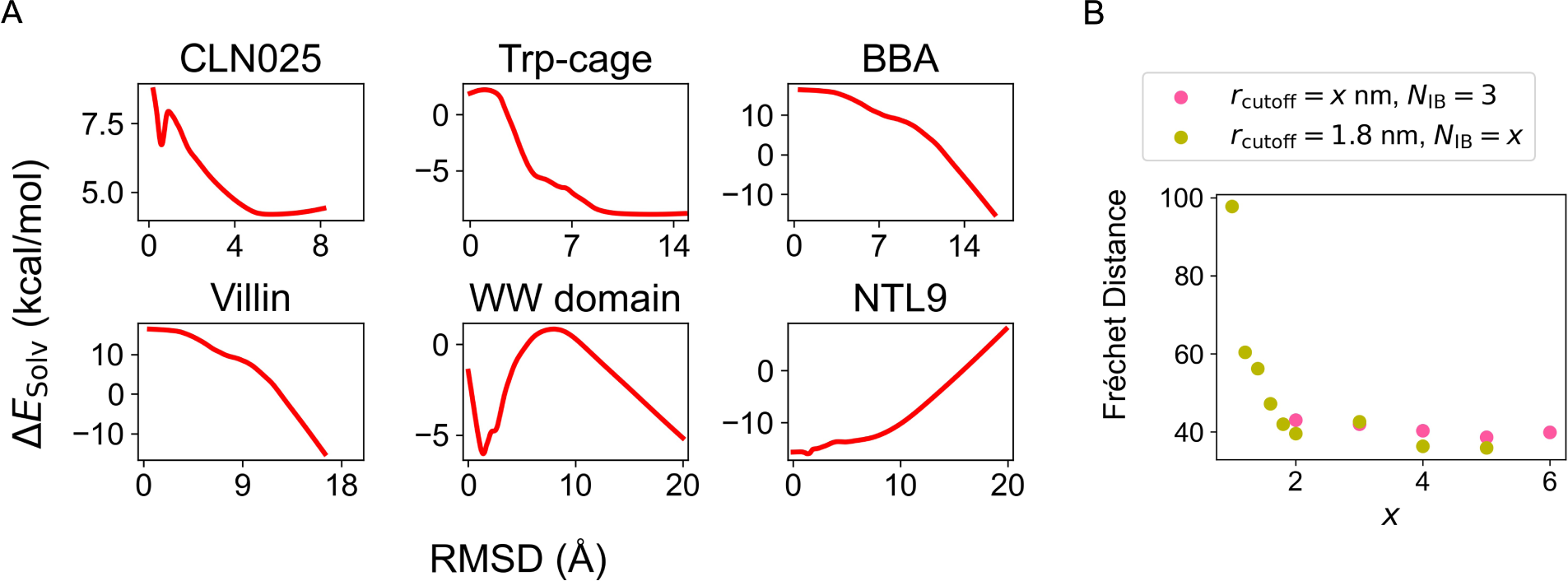
The solvation free energy learned from explicit solvent simulations by SchNet improves over the implicit solvent model, GBn2. (A) Average solvation free energy difference between the optimized SchNet and GBn2 models, i.e., Δ*E*_Solv_ = *E*_SchNet_*−E*_GBn2_, as a function of RMSD to the folded structures. We averaged over the configurations from explicit solvent simulations to compute Δ*E*_Solv_ at a given RMSD value and applied the LOWESS^92^ algorithm to smooth the curves. (B) The performance of the SchNet models optimized with potential contrasting at various hyperparameters. The performances are evaluated using the average discrete Fŕechet distance between atomistic and SchNet free energy profiles as a function of RMSD across all proteins. The SchNet free energy profiles used to compute these distances are shown in Figures S3 to S5.

### Pre-training for robust parameter optimization

We further explored the role of the hyperparameters, *r*_cut_ and *N*_IB_, on the accuracy of models optimized by potential contrasting. To assess the performance of the SchNet implicit solvent models, we computed the discrete Fŕechet distance^91^ between their free energy profiles and those from the explicit solvent simulations. The discrete Fŕechet distance is a widely used similarity measure between two arbitrary curves.

For all the hyperparameter combinations we examined, the sum of these distances across all proteins in the training set is presented in Figure 6B. For reference, the sum of the discrete Fŕechet distances between the explicit solvent and GBn2 free energy profiles for all proteins is 130.98. This metric suggests that all SchNet models outperform GBn2 in reproducing explicit solvent simulations. Consistent with the conclusions drawn from training SchNet to fit *E*_GBn2_, we observed that increasing *r*_cut_ leads to more accurate implicit solvent models. However, maintaining *r*_cut_ at 1.8 nm while varying the number of interaction blocks from *N*_IB_ = 3 does not significantly affect accuracy.

We observed that a pre-training procedure is essential for robust parameter optimization using potential contrasting. In this procedure, we initialize the SchNet parameters using those obtained from training the model to fit *E*_GBn2_. Additionally, we keep the parameters within the interaction blocks constant during the training process with potential contrasting. Without the initial pre-training, the free energy profiles depicted in Figure S6 indicate that the resulting model performs poorly, failing to replicate the results from explicit solvent simulations for all six proteins. Similarly, as shown in Figure S7, not holding the parameters within the interaction blocks constant also results in poor performance of the SchNet model.

Furthermore, we have discovered that pre-training enables stable ML-MD simulations. These simulations combine forces derived from the gas-phase CHARMM force field with those from the SchNet solvation free energy for the dynamic evolution of the equation of motion. For instance, we successfully conducted hundreds of nanoseconds of simulation for smaller proteins, such as CLN025 and Trp-cage, using SchNet models optimized with pretraining. Larger proteins impose greater demands on SchNet hyperparameters, but stable ML-MD simulations can still be performed with larger values for *N*_IB_. Conversely, SchNet models optimized without pre-training do not support long timescale simulations, regardless of the hyperparameter configuration.

It’s noteworthy that optimization with or without pre-training yielded comparable values of the loss function, indicating that it is an inadequate indicator of SchNet model performance. Upon examining the predicted solvation free energies *E*_SchNet_, it becomes evident that our pre-training scheme prevents SchNet from assigning unphysical energies (see Figures S8 and S9).

Our pre-training procedure is closely related to transfer learning, which involves transferring the knowledge a model has acquired from one task to another.^93^ Transfer learning is a widely employed technique to enhance the accuracy of machine learning models across a diverse range of problems.^94–96^

### SchNet enables highly accurate ML-MD simulations

We conducted hybrid ML-MD simulations to further assess the quality of the SchNet models trained using potential contrasting. Such simulations have the potential to yield more accurate free energy estimations by directly sampling configurations using the SchNet solvation free energy. In our investigation, we focused on a computationally more efficient SchNet architecture featuring three interaction blocks (*N*_IB_ = 3) and a cutoff distance of *r*_cut_ = 1.8 nm to define nearest neighbors. The free energy profiles computed using our re-weighting scheme indicate that this setup delivers accurate results, particularly for small proteins.

We performed 260 and 600 ns of RMSD-biased umbrella sampling simulations for CLN025 and Trp-cage, respectively. Details of these simulations can be found in the *Methods* section and the Supporting Information. Simulating larger systems with ML implicit solvents is computationally demanding and not attempted. Additional details regarding the free energy calculation methods are provided in the Supporting Information.

We observed that the free energy profiles computed from ML-MD simulations (Figure 7A) closely aligned with the re-weighted results (Figure S10) and exhibited strong agreement with the explicit solvent results. Additionally, the free energy profiles computed as a function of the radius of gyration (Rg) of all backbone heavy atoms from ML-MD simulations matched those from explicit solvent simulations (Figure S11A). These simulation outcomes affirm the accuracy of the derived SchNet model and validate the re-weighting scheme as a valuable tool for generating rapid and efficient approximations of free energy profiles.

**Figure 7:**
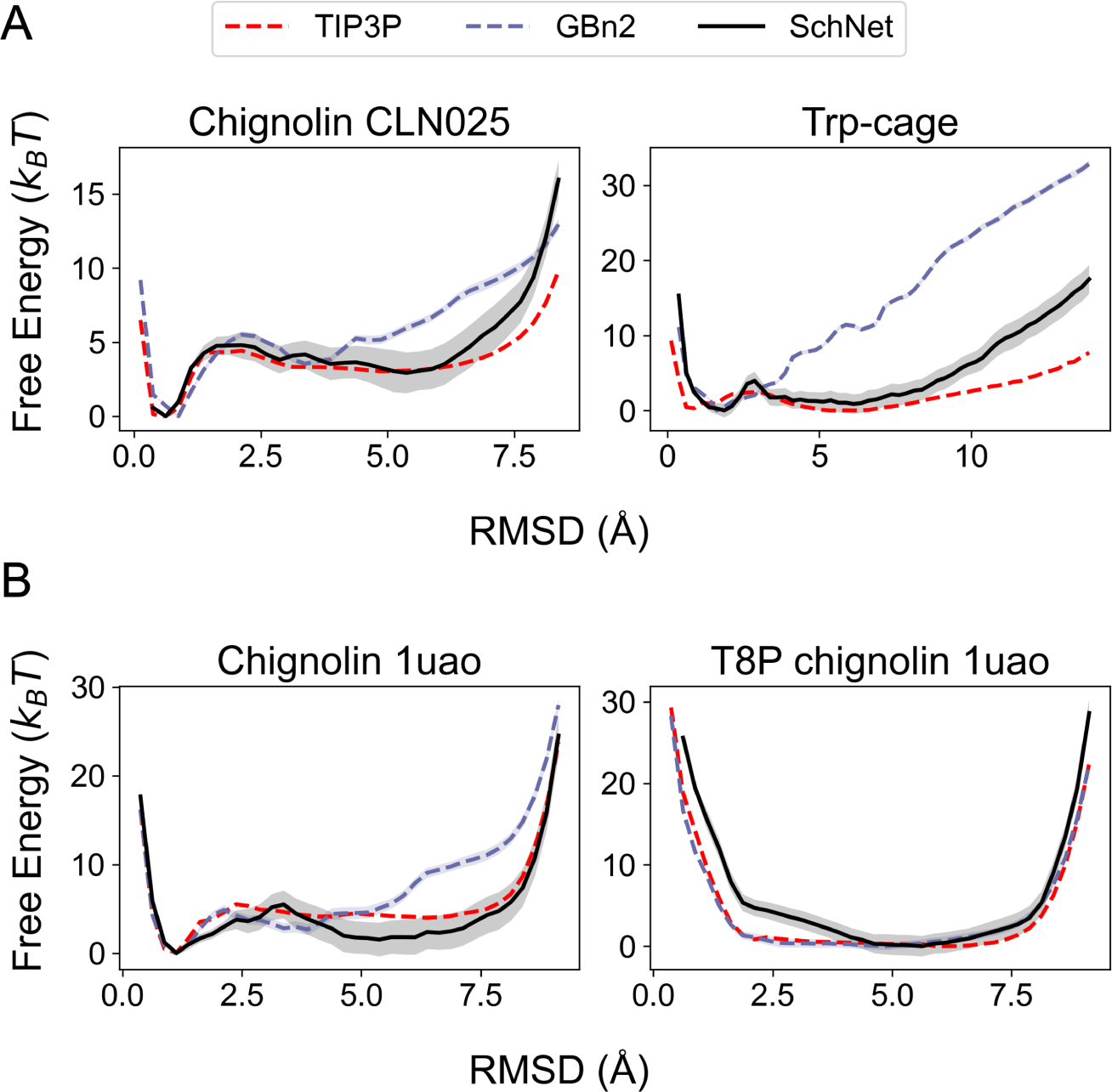
Comparison between the free energy profiles computed from explicit solvent simulations (TIP3P), implicit solvent simulations (GBn2), and the optimized SchNet model. Results for the SchNet model were determined using ML-MD umbrella simulations. (A) Free energy profiles for chignolin CLN025 and Trp-cage. TIP3P and GBn2 results are identical to those shown in Figure 5. (B) Free energy profiles for chignolin 1uao and a T8P mutant. TIP3P and GBn2 free energy profiles were computed from umbrella simulations. See the Supporting Information and Tables S4 to S5 for simulation details.

### ML-MD simulations show transferability of SchNet

We have demonstrated that a single SchNet model, trained using the potential contrasting method, accurately captures the conformational distribution of all six proteins in the training set (Figure 5). This remarkable achievement underscores the potential of GNNs as attractive alternatives for generating transferable representations of the solvation free energy. Next, we delve into assessing the transferability of the SchNet model by applying it to three proteins outside of the training set.

We first considered two variants of chignolin CLN025,^97^ denoted as 1uao and T8P. 1uao contains glycine residues at both termini, in contrast to tyrosine in the CLN025 sequence. ^98^ T8P, on the other hand, features a threonine-to-proline mutation at position 8 of 1uao, which discourages the native hairpin conformation. ^99^ These variants share 80% and 70% sequence identity, respectively, with CLN025.

We calculated free energy profiles through simulations using SchNet, GBn2, and TIP3P explicit solvent. Detailed simulation information can be found in the *Methods* section and the Supporting Information. As depicted in Figure 7B, for the 1uao system, SchNet once again outperforms GBn2 in predicting the relative stability of native and unfolded states. However, for the T8P mutant, SchNet’s performance is slightly inferior to that of GBn2, although it still accurately predicts the impact of the mutation, namely, the destabilization of the *β*-hairpin structure observed in 1uao. Free energy profiles as a function of Rg support these conclusions (see Figure S11B).

We further evaluated the SchNet model for an intrinsically disordered peptide (IDP), drastically different from the fast-folding proteins in our training set. The IDP corresponds to a structurally-active, 10-mer fragment of the C-Jun amino-terminal kinase-interacting protein 1 (JIP1), and has been studied extensively by Ojaghlou et al. ^100^ with explicit solvent simulations. It differs significantly from CLN025 with a sequence identity of 30%.

Free energy profiles as a function of Rg (Figure 8A) suggest that similar to GBn2, the SchNet model over-stabilizes the collapsed state. However, from the two-dimensional free energy profiles computed using the time-lagged independent component analysis (TICA) coordinates, we found the SchNet model correctly captures the three basins of the compact states explored in explicit solvent simulations (Figure 8B-C). Therefore, SchNet successfully predicted the right conformations, although it provided inaccurate free energies. Increasing the training set size, especially with the addition of disordered proteins, could further improve its transferability.

**Figure 8:**
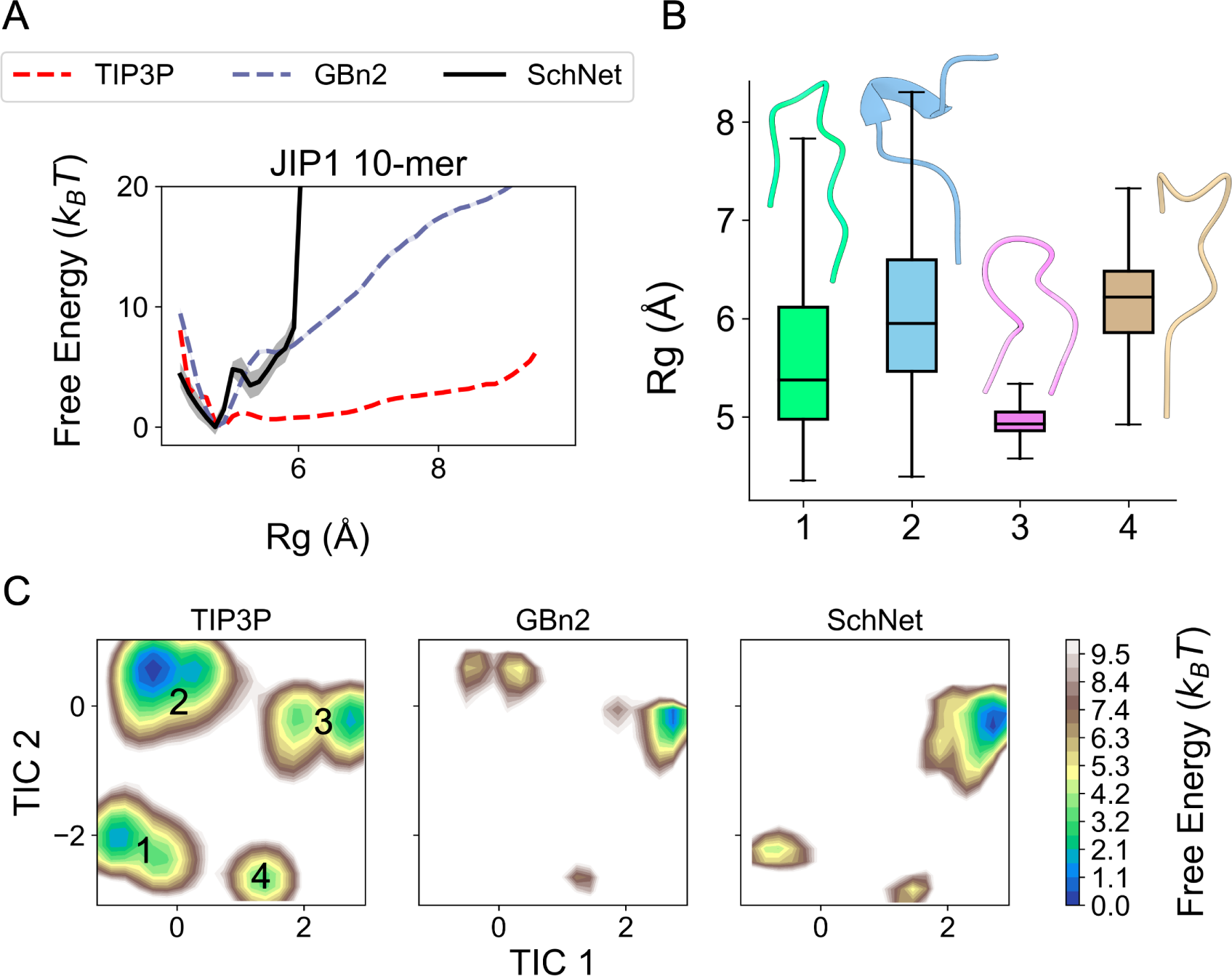
Comparing one- and two-dimensional free energy profiles between explicit solvent (denoted as TIP3P), GBn2 and the optimized SchNet model for the JIP1 10-mer. All explicit solvent results were generated by Ojaghlou et al. ^100^, while results for GBn2 and the SchNet model were generated from umbrella simulations. (A) Free energy profiles with respect to Rg. (B) Rg box plots and representative structures for each basin in the two-dimensional free energy profiles shown in part C. (C) Two-dimensional free energy profiles generated through TICA dimensionality reduction.^101–103^ The procedure used by Ojaghlou et al. ^100^ was followed (see Supporting Information for more detail).

## Conclusions and Discussion

We have introduced a comprehensive strategy for parameterizing GNN implicit solvent models from explicit solvent simulation data with the potential contrasting method. Several methodological advancements are crucial for enhancing the performance of the resulting models in describing the distribution of the entire configurational ensemble. These include a pre-training strategy for initializing parameters and a re-weighting scheme that facilitates efficient evaluation of hyperparameter combinations. Our results demonstrate that optimized SchNet models can accurately replicate the free energy profiles obtained from explicit solvent simulations for all six proteins in the training set. Simultaneously achieving such results for all six proteins represents a significant achievement, showcasing the model’s transferability. Furthermore, we have shown that the optimized SchNet model demonstrates reasonable transferability outside the training set.

When used with our re-weighting scheme, the parameterized SchNet model can offer immediate practical utility by delivering more precise solvation free energies than existing implicit solvent models. As an illustration, one can contemplate conducting simulations with GBn2 and subsequently applying the SchNet model to re-weight the obtained configurations, thereby bringing more accurate estimations of statistical properties.

However, ML-MD simulations using our SchNet implicit solvent model are considerably more computationally intensive than those utilizing the TIP3P explicit solvent. Table 1 compares the simulation speeds for CLN025, Trp-cage, and BBA in vacuum, explicit solvent, GBn2, and our SchNet implicit solvent model. Simulations with the SchNet implicit solvent (*N*_IB_ = 3, *r*_cut_ = 1.8 nm) are approximately 10 to 15 times slower than those in explicit solvent. These sluggish simulation speeds hinder effective conformational sampling for large proteins. We encountered several technical challenges that impeded more efficient simulations with our SchNet implicit solvent model, which we expect to be resolved in the near future. Firstly, we faced software dependency issues that prevented us from running simulations on the GPU. Secondly, our GNN implicit solvent implementation relies on slow communication between two software packages, PyTorch^104^ and OpenMM.^105^ The native integration of GNN models into MD simulation packages could circumvent this communication bottleneck and has the potential to significantly accelerate the simulations.

**Table 1:** Simulation speed for CLN025, Trp-cage, and BBA with different solvent models. Simulations for each protein were run on an Intel Xeon Platinum 8260 (4, 12, and 48 cores for CLN025, Trp-cage, and BBA, respectively) processor, and all simulation parameters are shown in Table S6.

| **System** | **CLN025** | **Trp-cage** | **BBA** |
| --- | --- | --- | --- |
| Vacuum (ns/day) | 103 | 80.3 | 89.1 |
| GBn2 (ns/day) | 51.2 | 43.1 | 51.8 |
| TIP3P (ns/day) | 32.4 | 35.0 | 22.1 |
| SchNet (ns/day) | 3.62 | 3.05 | 1.50<sup>a</sup> |

<sup>a</sup> Unstable with 5 fs time step used here, a 0.5 fs time step is required for ns simulations.

While our work primarily focused on parameterizing transferable implicit solvent models, the methodologies outlined here have broad applicability. They can be readily applied to developing lower-resolution CG models with fewer particles per amino acid. Utilizing GNNs as a flexible architecture for representing CG force fields means that accuracy does not have to be constrained by the approximations often required to derive analytical expressions for the PMF. Moreover, using potential contrasting enables robust parameterization of GNNs to replicate the conformational distribution observed in atomistic simulations. The combination of these two techniques has the potential to revolutionize coarse-graining practices, paving the way for the creation of transferable CG force fields that can rival the accuracy of explicit solvent atomistic simulations.

## Methods

### Detailed implementation of SchNet

As shown in Figure 1D, the SchNet architecture^52^ consists of three parts: an embedding layer (with learnable parameters ***θ*_0_**) for featurization, message-passing layers (with learnable parameters ***θ*_F_**) for feature updates, and a feed-forward NN (with learnable parameters ***θ*_E_**) for energy prediction. The embedding layer assigns a feature vector of *k* learnable parameters, 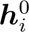, to the *i*th atom based on its atom type *t_i_*. Each message-passing layer, i.e., interaction block, further updates the feature vector of each atom by aggregating information from neighbors. Updates use continuous filters *W* learned from the distances between pairs of neighboring atoms. These filters are scaled by the distance between neighbors *i* and *j*, *r_ji_*, using a Behler-style cosine cutoff function *c* (*r_ji_*). The layers that compute *W* are shown in orange in Figure 1D. After *N*_IB_ total interaction blocks, the updated featurizations are inputted into the energy-predicting NN, which computes the total potential energy for the protein as a sum of individual contributions from each atom. Overall, the set of learnable parameters for SchNet is ***θ*** = *{****θ*_0_**, ***θ*_F_**, ***θ*_E_***}*. With the hyperparameters used to produce our optimal implicit solvent model (detailed in the Supporting Information and Table S7 - S8), each atom type is assigned *k* = 32 learnable parameters in the embedding layer, each interaction block contains 5,248 learnable parameters, and the energy-predicting NN contains 545 learnable parameters.

We mostly followed the implementation of SchNet from the PyTorch Geometric package,^106^ with only one change. Specifically, instead of featurizing each atom based on the atomic numbers, we used the atom types defined by the CHARMM force field. For example, we apply numeric labels to the atom type strings (outside of SchNet) and then use the embedding layer to assign ***h*_0_** to the numeric labels. As such, encoding atom types does not require additional layers. Like atomic numbers and partial charges, CHARMM atom types are available *a priori*. Since these atom types are defined based on neighboring atoms, the intention of this change was to speed convergence towards optimal parameters ***θ^∗^***.

### GNN optimization with potential contrasting

Potential contrasting builds upon the noise-contrastive estimation method^86^ to parameterize force fields. While it shares similarities with the maximum likelihood method, instead of directly maximizing the probability defined by the Boltzmann distribution of the force field on training data (i.e., configurations produced from atomistic simulations), the potential contrasting method optimizes force field parameters to best differentiate data configurations from noise configurations. The use of noise configurations circumvents the need to evaluate the partition function of the force field, thus significantly reducing the computational cost of optimization by eliminating the need for iterative sampling. The potential contrasting objective function, the loss function we minimize when parameterizing a force field, is detailed in the Supporting Information.

We generated noise configurations for each protein by conducting umbrella simulations employing the GBn2 implicit solvent model.^76^ These simulations involved applying biases on the RMSD from the folded structure, starting from 0.0 Å and incrementing by intervals of 0.5 Å up to their maximum RMSD, for each protein. Each simulation ran for a minimum duration of 155 ns, and configurations were sampled at intervals of 40, 50, or 200 picoseconds during the simulations (see Table S2). We merged the configurations obtained from different umbrella windows using free energy biases computed via MBAR.^107^ Further information on these umbrella simulations and the construction of the noise distribution are presented later in this section and the Supporting Information.

We conducted an initial pre-training phase before training SchNet with configurations from explicit solvent simulations using the potential contrasting method.^25^ This pre-training served two primary purposes: first, to assess SchNet’s ability to accurately predict the rather complex function representing the GBn2 solvation free energy, denoted as *E*_GBn2_, for all the proteins in the training set; and second, to learn the optimal parameters 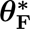 within the message passing layers. Consequently, the framework for performing feature updates became fully optimized through pre-training. The set of parameters acquired during this pre-training phase is denoted as 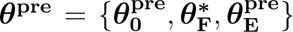. This pre-training process can be seen as imparting SchNet with the physical insights offered by the GBn2 implicit solvent model. As indicated in the text and illustrated in Figures S6, S7, and S9, in the absence of pre-training, potential contrasting, on its own, cannot ensure the satisfactory performance of the resulting SchNet models.

When training with potential contrasting, SchNet parameters were initialized as ***θ*^pre^**. In the subsequent optimization steps, only 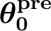 and 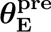 were permitted to change, while 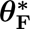 remained fixed. This approach effectively transferred the “knowledge” acquired by the message-passing layers during pre-training.

Training concluded after 120 epochs, corresponding to the number of times the loss function has been evaluated over the entire training set, for pre-training, and after 30 epochs for potential contrasting. This termination criterion ensured that both loss functions reached a plateau. The precise expressions of the loss functions are available in the Supporting Information. Both pre-training and potential contrasting employed the Adam optimizer.^108^ Initial learning rates of 1 *×* 10*^−^*^3^ and 1 *×* 10*^−^*^4^ were reduced by a factor of 10 every 15 and 10 epochs during pre-training and potential contrasting, respectively. During pre-training, the learning rate was reset to 1 *×* 10*^−^*^3^ after epoch 60 and subsequently decreased by a factor of 10 every 15 epochs until training completion.

### Re-weighting configurations to estimate free energy profiles

Evaluating the accuracy of our SchNet implicit solvent model can be challenging, primarily because we lack precise values for the solvation free energy that the model aims to reproduce. Conducting extensive sampling through ML-MD simulations enables the creation of low-dimensional probability distributions of selected collective variables, which can then be compared to explicit solvent simulation results. While this evaluation technique is reliable, it is often computationally demanding. To address this, we propose a straightforward reweighting scheme using the free energy perturbation method^78^ to swiftly generate reasonably accurate estimates of our SchNet implicit solvent models’ performance.

We introduce a weighting factor, denoted as 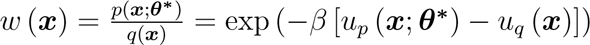, for each noise configuration. Here, *u_q_* (***x***) represents the energy function of the noise distribution, while *u_p_* (***x***; ***θ^∗^***) corresponds to the energy defined by the optimized SchNet. These weights are subsequently normalized so that max *w* (***x***) = 1. Using the noise configurations and their corresponding weighting factors, we construct a weighted histogram with *L* RMSD bins, where we choose *L* to ensure that the RMSD width of each bin is 0.25 Å.

Within each of the *n_l_* configurations present in the *l*th bin, we compute the free energy, denoted as 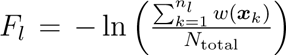. To ensure that the set of free energies for all *L* bins, 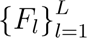, aligns with a physically meaningful scale, we uniformly shift it to ensure that 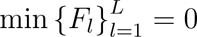 in units of *k_B_T*. Recognizing that physically meaningful free energy should exhibit smoothness, we further apply the LOWESS^92^ smoothing scheme to 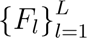. This process yields a reasonably accurate estimate of the free energy as a function of RMSD.

Additionally, we compute standard deviations for each re-weighted free energy by employing the block bootstrapping^109^ method on the noise configurations and their corresponding energies. The block size was set as the number of configurations collected from 2 ns of simulation. Ten bootstrapping samples were generated to calculate ten estimates, and their average and standard deviation are used as the free energy profile and the error bars in the figures.

### Molecular dynamics simulation details

RMSD-biased umbrella simulations were conducted for all proteins, with RMSD computed based on the heavy atoms in the peptide backbone in relation to the folded structure of each protein. These simulations were carried out using either the GBn2 or the SchNet implicit solvent model, with either the CHARMM^87–89^ or AMBER^110^ gas-phase force field. The Langevin middle integrator was used for all simulations.^111^ All umbrella simulations were performed with the OpenMM package^105^ (version 8.0 beta). For simulations employing the SchNet model, the OpenMM-Torch package^112^ (version 1.0 beta) was also utilized. Further simulation details can be found in the Supporting Information.

Inputs were generated using CHARMM-GUI.^41,113,114^ Trajectory processing and the calculation of RMSD and Rg were accomplished using the MDTraj^115^ and pytraj^116^ packages. Aditional analyses were performed with the aid of the statsmodels^92,117^ (version 0.14.0) and scipy^118,119^ packages. All structure visualization was performed using ChimeraX.^120^

Configurations generated from the umbrella simulations were utilized to calculate the free energy as a function of RMSD or Rg. We employed the MBAR^107^ and UWHAM^121^ methods for these calculations, utilizing the FastMBAR package.^122^ To assess the reliability of our results, we determined standard deviations for the free energy profiles through the block bootstrapping technique.^109^ Following the same procedure used for the re-weighted free energy profiles, the block size was set as the number of configurations collected from 2 ns of simulation, and ten bootstrap samples were created to compute average and standard deviation of the free energy profile. For more detailed information regarding the free energy calculations, please refer to the Supporting Information.

## Acknowledgement

This work was supported by the National Institutes of Health (Grant R35GM133580). We acknowledge the MIT Supercloud and Lincoln Laboratory Supercomputing Center for providing high-performance computing resources. We thank D. E. Shaw Research for sharing their atomistic simulation data. J.A. was the recipient of the MIT Dean of Science Fellowship and the National Science Foundation Graduate Research Fellowship.

