## Supporting Information for "Transferable Coarse Graining via Contrastive Learning of Graph Neural Networks"

#### Contents

**Selecting data samples from explicit solvent MD simulations** **S-1**

**Umbrella sampling simulation set-up** **S-2**

Umbrella simulations with GBn2 to generate noise configurations . . . . . . . . . . S-3

MD simulations for proteins outside of the training set . . . . . . . . . . . . . . . . S-4

ML-MD simulations using the SchNet implicit solvent . . . . . . . . . . . . . . . . . S-6

Benchmarking simulations . . . . . . . . . . . . . . . . . . . . . . . . . . . . . . . S-7

Software versions & hardware . . . . . . . . . . . . . . . . . . . . . . . . . . . . . S-7

**Pre-training & potential contrasting set-up** **S-8**

Loss functions . . . . . . . . . . . . . . . . . . . . . . . . . . . . . . . . . . . . . S-8

Hyperparameters . . . . . . . . . . . . . . . . . . . . . . . . . . . . . . . . . . . S-10

Software versions & hardware . . . . . . . . . . . . . . . . . . . . . . . . . . . . . S-11

**Noise potential & free energy profile calculations** **S-12**

Constructing the noise potential using FastMBAR . . . . . . . . . . . . . . . . . . . S-13

Free energy profile calculations . . . . . . . . . . . . . . . . . . . . . . . . . . . . S-14

Calculating free energy profiles from umbrella simulations using FastMBAR . S-14

Calculating free energy profiles from unbiased explicit solvent simulations . . S-15

TICA free energy profiles . . . . . . . . . . . . . . . . . . . . . . . . . . . . . . S-16

**Figures** **S-17**

Table S1: Interval, RMSD cutoff, and numbers of removed configurations for each protein for the selection of data samples. The RMSD removed row indicates the number of frames removed based on the selected RMSD cutoff, while the front and end removed rows indicate the number of configurations discarded from the beginning and the end of each ensemble. This selection process resulted in 100,000 data configurations for each protein. See text section *Selecting data samples from explicit solvent MD simulations* for further details.

| **System** | **CLN025** | **Trp-cage** | **BBA** | **Villin** | **WW domain** | **NTL9** |
|---|---|---|---|---|---|---|
| Interval[a] | 5 | 5, 2 | 8, 2 | 5 | 25, 2 | 67, 2 |
| RMSD cutoff (nm)[b] | 1 | 1.5 | 1.75 | 2 | | |
| RMSD removed | 0 | | | | 88 | 4 |
| Front removed | 3,000 | 4,400 | 1,250 | 12,000 | 13,550 | 10,500 |
| End removed | 3,000 | 4,400 | 1,250 | 12,000 | 13,562 | 10,596 |

[a] If listed, a 2nd interval was used to select frames after RMSD-based, front, and end configuration removal.
[b] RMSD cutoff selected based on RMSD ranges from umbrella sampling MD (Table S2).

### Selecting data samples from explicit solvent MD simulations

Atomistic TIP3P[S1] explicit solvent molecular dynamics (MD) simulations for chignolin CLN025 (CLN025), Trp-cage, BBA, Villin, WW domain, and NTL9 were conducted by Lindorff-Larsen et al.[S2] From each of the six ensembles, we selected 100,000 configurations as data samples for training. The process for selecting data samples for each protein was as follows: first configurations were selected at a specific interval, next a small number of configurations beyond a specific peptide backbone heavy atom root-mean-squared-deviation (RMSD) distance cutoff from the folded structure $\boldsymbol{x}_\circ$ were removed, and lastly a number of configurations were removed from the beginning and the end of the ensembles to ensure exactly 100,000 configurations for all proteins. Configurations beyond the RMSD cutoff were removed as these fell outside of the range covered by the noise configurations. Few configurations fell beyond RMSD the cutoff, if at all. All intervals, RMSD cutoffs, and beginning (front)- and end-of-ensemble removed frames are detailed in Table S1.

$k$-means clustering (using the `cluster.kmeans` function of the pytraj package[S3]) was used to identify $\boldsymbol{x}_\circ$ for each protein. The centroid frame of the most populated of $k = 10$ computed clusters was selected as $\boldsymbol{x}_\circ$ for each protein. A similar process was used to identify $\boldsymbol{x}_\circ$ in the original study by Lindorff-Larsen et al.[S2]. The structures we identified as $\boldsymbol{x}_\circ$ match those identified by Lindorff-Larsen et al.[S2]

### Umbrella sampling simulation set-up

RMSD-biased umbrella sampling MD simulations were conducted for each protein in the training set using the GBn2 implicit solvent model model.[S4] We also performed similar simulations using the trained SchNet implicit solvent model.

The RMSD was computed in reference to the folded structures, $\boldsymbol{x}_\circ$, identified using a procedure detailed in the previous section *Selecting data samples from explicit solvent MD simulations*. Only peptide backbone heavy atoms were used to compute the RMSD. A harmonic RMSD-biasing potential

$$u^i_{\text{bias}}\left(\boldsymbol{x}\right) = \frac{1}{2}k^i_{\text{RMSD}}\left[\text{RMSD}\left(\boldsymbol{x}, \boldsymbol{x}_\circ\right) - \text{RMSD}^\circ_i\right]^2 \quad \text{(S1)}$$

was applied to the $i$th umbrella window. A total of $M$ windows were sampled for each protein. The RMSD centers, $\text{RMSD}^\circ_i$, for each protein were evenly spaced at an interval of 0.05 nm within the RMSD range indicated in the tables that follow. RMSD ranges were selected to cover the full configurational space of each protein (Figure S12).

For all umbrella sampling simulations (with exceptions detailed later), a force constant $k^i_{\text{RMSD}} = 15,000$ kJ/mol/nm$^2$ was applied to the first two ($i = 1$ and $i = 2$) and the last two ($i = M - 1$ and $i = M$) windows within the indicated RMSD ranges. $k^i_{\text{RMSD}} = 12,500$ kJ/mol/nm$^2$ was applied to the next two ($i = 3$ and $i = 4$) and the next-to-last two ($i = M-3$ and $i = M-2$) windows within the indicated ranges. For all other windows, $k^i_{\text{RMSD}} = 10,000$ kJ/mol/nm$^2$ was applied.

Minimization and equilibration were performed prior to all production simulations. Min-

Table S2: Parameters for RMSD-biased umbrella sampling MD simulations for proteins within the training set.

| **System** | **CLN025** | **Trp-cage** | **BBA** | **Villin** | **WW domain** | **NTL9** |
|---|---|---|---|---|---|---|
| Solv. model | GB-neck2 (GBn2)[S4] | | | | | |
| Force fields[a] | CHARMM36 (C36)[S6] | | | C36m[S7] C36[S8] | C36[S6] | C36m[S7] C36[S8] |
| Integrator | Langevin Middle[S9] | | | | | |
| Temperature (k)[b] | 340 | 290 | 325 | 360 | | 355 |
| Friction $\gamma$ ($ps^{-1}$) | 1 | | | | | |
| Constraints | H bonds[c] | | | | | |
| H-mass (amu) | 1 | | | | | |
| Nonbond scheme | No cutoff | | | | | |
| Time step (fs) | 2.5 | | | | | |
| Minimize steps | 50,000 | | | | | |
| Equilibrate steps | 1,200,000 | | | | | |
| Traj. length (ns)[d] | 1,060 | 1,392 | 225 | 155 | 284 | 276.25 |
| Save interval (ps) | 200 | | 40 | 50 | | |
| RMSD (nm) | 0 - 0.95 | 0 - 1.45 | 0 - 1.75 | 0 - 1.95 | | |
| $M$[e] | 20 | 30 | 36 | 40 | | |

[a] CHARMM36m[S7] (C36m) and C36 for nonstandard amino acids[S8] were used to parameterize norleucine residues in villin and NTL9. C36 for nonstandard amino acids was validated for use with C36m.[S8]
[b] Same temperatures as Lindorff-Larsen et al.[S2] simulations.
[c] H-X bond lengths constrained.
[d] Length of each umbrella window trajectory (traj.).
[e] Total # of umbrella windows.

imization was performed using the `minimizeEnergy` method of the `Simulation` class in OpenMM.[S5] Equilibration was performed by iteratively increasing $k^i_{\mathrm{RMSD}}$ to its production value. For each protein, the tables that follow detail the total number of minimization and equilibration steps and time steps, respectively.

#### Umbrella simulations with GBn2 to generate noise configurations

RMSD-biased umbrella sampling MD simulations were conducted to generate noise samples for the following training set proteins: CLN025, Trp-cage, BBA, Villin, WW domain, and

Table S3: Selection ranges and intervals of noise sample configurations. The process for selecting noise sample configurations was as follows: first configurations were removed from the beginning of each umbrella window, next the remaining configurations were concatenated into a single umbrella ensemble, and lastly configurations were selected from the umbrella ensemble at a specific interval. Ranges of selected configurations are shown for each umbrella window, while the interval at which frames were selected are shown for the entire umbrella ensemble. An additional frame was occasionally removed from some windows to ensure 100,000 configurations for each protein.

| **System** | **CLN025** | **Trp-cage** | **BBA** | **Villin** | **WW domain** | **NTL9** |
|---|---|---|---|---|---|---|
| Range(s) | [300:] | [293:], [294:][a] | [69:], [70:][b] | [600:] | [680:] | [525:] |
| Interval | 1 | 2 | | 1 | 2 | |

[a] An additional frame was removed from windows in range 1 - 1.45 nm.
[b] An additional frame was removed from windows in range 1 - 1.75 nm.

NTL9. Table S2 details the simulation parameters, number of minimization steps, number of equilibration time steps, RMSD ranges, and total number of umbrella windows $M$ used for these umbrella sampling simulations. For each protein, the interval and range of noise sample configurations selected from each umbrella ensemble are detailed in Table S3. From the resultant umbrella ensembles, 100,000 configurations for each protein were used as noise samples for training with the potential contrasting[S10] method.

Note that for CLN025 and Trp-cage, each umbrella trajectory is of significantly longer timescale than for the remaining proteins in the training set (Table S2). While it was initially unknown what timescale would be necessary to achieve convergence of the free energy profile with respect to RMSD for each protein, later analysis revealed that such long timescales were unnecessary. As such, trajectories for the remaining proteins are of shorter timescale and save configurations at a higher frequency. Since our configurations sufficiently sample the RMSD phase space (Figure S12), it is unlikely that timescale differences impact the training with potential contrasting. Free energy profile convergence for each protein is demonstrated in Figure S13, while the process for computing these free energy profiles is detailed in section *Noise potential & free energy profile calculations*.

Table S4: Parameters used for TIP3P and GBn2 RMSD-biased umbrella sampling MD simulations for proteins outside of the training set.

| **System** | **1uao** | | **T8P** | | **JIP1 10-mer** |
|---|---|---|---|---|---|
| Solv. model | TIP3P[S1] | GBn2[S4] | TIP3P[S1] | GBn2[S4] | GBn2[S4] |
| Force fields | C36[S6] | | | | ff14SB[S11] |
| Integrator | Langevin Middle[S9] | | | | |
| Temperature (K) | 315[a] | | 420[b] | | 300[c] |
| Friction $\gamma$ ($ps^{-1}$) | 1 | | | | 0.01 |
| Constraints | H bonds[d] | | | | All bonds |
| H-mass (amu) | 1 | | | | 4 |
| Nonbond scheme | PME | No cutoff | PME | No cutoff | No cutoff |
| Nonbond cutoff (nm) | 1.2 | - | 1.2 | - | - |
| Switch dist. (nm) | 1 | - | 1 | - | - |
| Solv. padding (nm) | 2.4 | - | 2.4 | - | - |
| Time step (fs) | 2.5 | | | | 5 |
| Minimize steps | 50,000 | | | | |
| Equilibrate steps | 1,200,000 | | | | 1,100,000 |
| Traj. length (ns)[e] | 62.5 | | | | 1,000 |
| Save interval (ps) | 25 | | | | 20 |
| RMSD (nm) | 0 - 0.95 | | | | |
| $M^f$ | 20 | | | | |

[a] Denaturation midpoint (melting temperature) for 1uao.
[b] Gly7-Pro8 *cis* to *trans* transition temperature for T8P 1uao mutant.
[c] Same temperature used by Ojaghlou et al.[S12]
[d] H-X bond lengths constrained.
[e] Length of each umbrella window traj.
[f] Total # of umbrella windows.

#### MD simulations for proteins outside of the training set

Table S4 details the simulation parameters, number of minimization steps, number of equilibration time steps, RMSD ranges, and total number of umbrella windows $M$ for umbrella sampling simulations used to generate atomistic, TIP3P explicit solvent and GBn2 implicit solvent configurations for the following proteins: chignolin 1uao (PDB ID: 1uao),[S13] a T8P mutant of chignolin 1uao that does not favor the native hairpin structure,[S14] and a structurally-active 10-mer of the C-Jun amino-terminal kinase-interacting protein 1 (JIP1).[S12] These proteins were used to evaluate the transferability of the SchNet implicit solvent model outside the training set. The explicit solvent configurations were generated by Ojaghlou

Table S5: Parameters used for RMSD-biased umbrella sampling ML-MD simulations using the SchNet implicit solvent. Parameters for CLN025, Trp-cage, 1uao chignolin, T8P 1uao chignolin, and the JIP1 10-mer are shown.

| **System** | **CLN025** | **Trp-cage** | **1uao** | **T8P** | **JIP1 10-mer** |
|---|---|---|---|---|---|
| Solv. model | SchNet implicit solvent | | | | |
| Force fields | C36[S6] | | | | ff14SB[S11] |
| Integrator | Langevin Middle[S9] | | | | |
| Temperature (K) | 340 | 290 | 315 | 420 | 300 |
| Friction $\gamma$ ($ps^{-1}$) | 1 | | 1 | | 0.01 |
| Constraints | All bonds | | All bonds | | |
| H-mass (amu) | 4 | | | | |
| Nonbond scheme | No cutoff | | | | |
| Time step (fs) | 5 | | | | |
| Minimize steps | 5,000 | | | | 1,000 |
| Equilibrate steps | 540,000 | 600,000 | 540,000 | | |
| Traj. length (ns)[a] | 13 | 20 | 13 | | 30 |
| Save interval (ps) | 25 | 20 | 25 | | 10 |
| RMSD (nm) | 0 - 0.95 | 0 - 1.45 | 0 - 0.95 | | |
| $M^b$ | 20 | 30 | 20 | | |

[a] Length of each umbrella window traj.
[b] Total # of umbrella windows.

et al.[S12] for the JIP1 10-mer and by new umbrella sampling MD simulations for 1uao and the T8P mutant. GBn2 configurations were also generated using umbrella sampling MD simulations for all three proteins. Since there are 18 individual structures associated with the PDB ID 1uao, the structure with the lowest gas-phase energy (the 15$^{\text{th}}$ structure) was used as $\boldsymbol{x}_\circ$. The same structure (albeit with the T8P mutation) was used as $\boldsymbol{x}_\circ$ for the T8P mutant. The T8P mutant was generated by using the `PDB Reader & Manipulator` tool in CHARMM-GUI[S15–S17] to mutate the selected 1uao PDB structure. While the JIP1 10-mer lacks a distinct folded state, the compact structure of Conformer 1 as identified by Ojaghlou et al.[S12] was used as $\boldsymbol{x}_\circ$. Free energy profile convergence for umbrella sampling simulations of these proteins is displayed in Figure S14.

#### ML-MD simulations using the SchNet implicit solvent

Table S5 details the simulation parameters, number of minimization steps, number of equilibration time steps, RMSD ranges, and total number of umbrella windows $M$ used for RMSD-biased umbrella sampling simulations using the SchNet implicit solvent. Umbrella sampling was conducted to enable the calculation of free energy profiles with respect to RMSD from $\boldsymbol{x}_\circ$ for comparison to the results from atomistic TIP3P and GBn2 simulations. Note that while our SchNet implicit solvent model was developed with the intention of using it with C36, ff14SB[S11] was used for the JIP1 10-mer since the explicit solvent configurations were generated by Ojaghlou et al.[S12] using ff14SB. Free energy profile convergence for these simulations is displayed in Figure S15.

#### Benchmarking simulations

Table S6 details the parameters of simulations used to evaluate the simulation speed of the SchNet implicit solvent. All simulations were run with a harmonic RMSD biasing potential (Eq. S1) centered at 0.0 nm from the folded structures. Compared with more extended conformations, folded structures produce more nearest neighbors per atom when defining the graph, creating a worst-case scenario for the SchNet implicit solvent model.

#### Software versions & hardware

MD simulations with TIP3P explicit solvent or GBn2 implicit solvent were conducted using OpenMM[S5] version (ver.) 7.7.0 on a single Nvidia Volta V100 GPU. ML-MD simulations with the SchNet implicit solvent model were conducted using OpenMM[S5] ver. 8.0.0 beta and OpenMM PyTorch plugin[S18] ver. 1.0 beta with PyTorch[S19] ver. 1.11.0 and PyTorch Cluster[S20] ver. 1.5.9. Due to software dependency limitations, ML-MD simulations were run on Intel Xeon CPUs, with 4 cores for CLN025, 12 cores for Trp-cage, 12 cores for 1uao, 8 cores for T8P 1uao, and 4 cores for the JIP1 10-mer. Trajecotry analyses such as RMSD

Table S6: Parameters for MD and ML-MD simulations used for benchmarking.

| **System** | **CLN025** | | **Trp-cage** | | **BBA** | |
|---|---|---|---|---|---|---|
| Solv. model | Vacuum / GBn2[S4]/ SchNet | TIP3P[S1] | Vacuum / GBn2[S4]/ SchNet | TIP3P[S1] | Vacuum / GBn2[S4]/ SchNet | TIP3P[S1] |
| Force fields | ff14SB[S11] | | | | | |
| Integrator | Langevin Middle[S9] | | | | | |
| Temperature (K) | 340 | | 290 | | 325 | |
| Friction $\gamma$ ($ps^{-1}$) | 1 | | 1 | | 0.01 | |
| Constraints | All bonds | | All bonds | | H Angles[a] | |
| H-mass (amu) | 4 | | | | | |
| Nonbond scheme | No cutoff | PME | No cutoff | PME | No cutoff | PME |
| Nonbond cutoff (nm) | - | 1.2 | - | 1.2 | - | 1.2 |
| Switch dist. (nm) | - | 1 | - | 1 | - | 1 |
| Solv. padding (nm) | - | 2.4 | - | 2.4 | - | 2.4 |
| Time step (fs) | 5 | | | | | |
| Minimize steps | 5,000 | | | | | |
| Equilibrate steps | 540,000 | | 600,000 | | 250,000 | |
| Traj. length (ns) | 12 | | 7 | | 0.875 | |
| Save interval (ps) | 25 | | 20 | | 0.5 | |
| RMSD center (nm) | 0.0 | | 0.0 | | 0.0 | |

[a] All bond lengths, and all H-X-H and H-O-X angles fixed

and $k$-means clustering were performed using both pytraj[S3] ver. 2.0.6 and MDtraj[S21] ver. 1.9.7.

### Pre-training & potential contrasting set-up

#### Loss functions

The root-mean-squared-error (RMSE) loss function was used for pre-training the SchNet[S22] model to fit $E_{\mathrm{GBn2}}$. This was implemented in PyTorch[S19] as the square root of the `MSELoss` class

$$\mathrm{RMSE} = \sqrt{\frac{\sum_{i=1}^{N_{\mathrm{set}}} \left[E^{i}_{\mathrm{GBn2}}\left(\boldsymbol{x}\right) - E^{i}_{\mathrm{SchNet}}\left(\boldsymbol{x};\boldsymbol{\theta}\right)\right]^2}{N_{set}}}, \tag{S2}$$

where $N_{\mathrm{set}}$ is the total number of configurations in the training / testing sets. The dataset used to pre-train SchNet was composed of 100,000 atomistic configurations and 100,000 GBn2 configurations per protein. These were the same configurations used to train SchNet with the potential contrasting method. The testing set held aside when pre-training SchNet was composed of 50,000 configurations. As such, for the training set $N_{\mathrm{set}} = 150,000$ and for the testing set $N_{\mathrm{set}} = 50,000$.

The objective function used in potential contrasting is defined as

$$\ell_{\mathrm{tot}}\left(\boldsymbol{\theta}, \{\boldsymbol{\Delta F}\}_{k=1}^{K}\right) = \sum_{k=1}^{K} \frac{1}{N_p^k} \Bigg\{ \sum_{i=1}^{N_p^k} \ln \left[ \frac{1}{1 + \nu_k \exp\left(-\beta \left[u_q^k\left(\boldsymbol{x}_p^{ki}\right) - u_p\left(\boldsymbol{x}_p^{ki};\boldsymbol{\theta}\right) + \boldsymbol{\Delta F}_k\right]\right)} \right] + \\ \sum_{i=1}^{N_q^k} \ln \left[ \frac{1}{1 + \nu_k^{-1} \exp\left(-\beta \left[u_p\left(\boldsymbol{x}_q^{ki};\boldsymbol{\theta}\right) - u_q^k\left(\boldsymbol{x}_q^{ki}\right) - \boldsymbol{\Delta F}_k\right]\right)} \right] \Bigg\}, \tag{S3}$$

where $k$ indices over proteins in the training set. $\boldsymbol{\Delta F}_k = \boldsymbol{F}_p^k - \boldsymbol{F}_q^k$ and $\nu_k = \frac{P(y=0)}{P(y=1)} = \frac{N_q}{N_p}$. $\boldsymbol{\Delta F}$ is a free parameter during training. When training SchNet with potential contrasting, the bias parameter of the final layer of the feed-forward NN is held constant to prevent it from counter-acting $\boldsymbol{\Delta F}$. $u_q^k\left(x\right)$ is the potential energy function corresponding with the noise distribution of the $k$th protein, while $u_p\left(\boldsymbol{x};\boldsymbol{\theta}\right) = U_{\mathrm{C36}}\left(x\right) + E_{\mathrm{SchNet}}\left(\boldsymbol{x};\boldsymbol{\theta}\right)$ is the potential energy corresponding with the learned CG distribution that approximates the atomistic distribution. $U_{\mathrm{C36}}$ is simply the potential energy computed from the CHARMM force field.[S6,S7] We used the same number of data samples $N_p^k$ = 100,000 and noise samples $N_q^k$ = 100,000. As such $\nu_k = 1$ for all proteins in our training set and will be omitted from all equations that follow. Equation S3 was implemented in PyTorch as the sum of individual binary cross entropy (BCE) with log-odds-ratios (logits) loss functions for each protein

$$\mathrm{BCE}\left(\boldsymbol{\theta}, \{\boldsymbol{\Delta F}\}_{k=1}^{K}\right) = \sum_{k=1}^{K} \left\{ \frac{1}{N_{tot}^{k}} \sum_{i=1}^{N_{tot}^{k}} y_{ki} \ln\left[\sigma\left(\mathrm{logit}\left(\boldsymbol{x}^{ki}\right)\right)\right] + \right.$$
$$\left. \left(1-y_{ki}\right) \ln\left[1-\sigma\left(\mathrm{logit}\left(\boldsymbol{x}^{ki}\right)\right)\right] \right\}, \quad \text{(S4)}$$

where $\sigma(z) = 1/\left[1+\exp(-z)\right]$ is the sigmoid function and $y$ is a label that indicates whether $\boldsymbol{x}^{ki}$ is a data sample ($y=1$) or noise sample ($y=0$). $N_{\mathrm{tot}}^{k} = N_{p}^{k} + N_{q}^{k}$ was the total number of configurations for the $k$th protein. Each individual BCE with logits loss term was implemented with the `BCEWithLogitsLoss` class in PyTorch. By inserting the logit of $\boldsymbol{x}_{ki}$ for our learned $p(\boldsymbol{x};\boldsymbol{\theta})$ and our noise $q_k(\boldsymbol{x})$ distributions,

$$\mathrm{logit}\left(\boldsymbol{x}^{ki}\right) = \ln\left(\frac{p\left(\boldsymbol{x}^{ki};\boldsymbol{\theta}\right)}{q_k\left(\boldsymbol{x}^{ki}\right)}\right) = \ln\left(\frac{\exp\left(-\beta\left[u_p\left(\boldsymbol{x}^{ki};\boldsymbol{\theta}\right) - \boldsymbol{F}_p^{k}\right]\right)}{\exp\left(-\beta\left[u_q^{k}\left(\boldsymbol{x}^{ki}\right) - \boldsymbol{F}_q^{k}\right]\right)}\right)$$
$$= -\beta\left[u_p\left(\boldsymbol{x}^{ki};\boldsymbol{\theta}\right) - u_q^{k}\left(\boldsymbol{x}^{ki}\right) - \boldsymbol{\Delta F}_k\right], \quad \text{(S5)}$$

into $\sigma(z)$, it is clear that equations S3 and S4 are effectively equivalent.

An L2 regularization term $\lambda\sum_{i=1}^{N_{\mathrm{parms}}}\boldsymbol{\theta}_i^2$ (where $N_{\mathrm{parms}}$ is the total number of trainable parameters in SchNet, $\boldsymbol{\theta}_i$ represents the $i$th trainable parameter, and $\lambda$ represents the regularization constant hyperparameter) was added to both loss functions (equations S2 and S4) during training to help prevent overfitting. This was implemented in PyTorch by specifying a `weight_decay` value within the `optim.Adam` class.

#### Hyperparameters

Table S7 details all explored SchNet architectures,[S22] while Table S8 details the training set-up hyperparameters (learning rates $\zeta$, L2 regularization constants $\lambda$, batch sizes, etc.). As discussed in the main text, the cutoff distance used for defining nearest neighbors ($r_{\mathrm{cut}}$) and the number of interaction blocks ($N_{\mathrm{IB}}$) are the most impactful hyperparameters for the

SchNet architecture. Increasing the embedding width only appeared to increase memory usage, and as such no rigorous evaluation of this hyperparameter was performed.

With all other optimal hyperparameters selected (Table S7 - S8), several SchNet implicit solvents with different values of the L2 regularization constant $\lambda$ were trained. While not found to affect re-weighted free energy profiles, $\lambda$ had a significant effect on the performance of the SchNet implicit solvent model in ML-MD simulations. Unlike when performing re-weighting, computing free energy profiles from ML-MD simulations involves entirely new configurations (i.e., not included in the training set). Since it is well-known that regularization can improve the ability of ML models to generalize to samples outside of the training set, this outsized role of $\lambda$ is not unexpected.

For models that we attempted to run ML-MD simulations with, the optimal value of $\lambda$ was selected by running umbrella simulations for CLN025 and computing free energy profiles with respect to RMSD from $\boldsymbol{x}_{\circ}$. For the hyperparameter combination used to generate the ML-MD simulation data discussed in the main text ($N_{\mathrm{IB}} = 3$, $r_{\mathrm{cut}} = 1.8$ nm), CLN025 free energy profiles computed from ML-MD for models with different values of $\lambda$ are shown in Figure S16. The most accurate model ($\lambda = 1 \times 10^{-6}$) was used for all simulations discussed in the main text. For a SchNet implicit solvent model trained with the same hyperparameters but without holding the interaction block parameters constant, free energy profiles are shown in Figure S17. Lastly, for a smaller SchNet model ($N_{\mathrm{IB}} = 2$, $r_{\mathrm{cut}} = 1.8$ nm), free energy profiles are shown in Figure S18. While one of these models was accurate for CLN025 ($\lambda = 1 \times 10^{-3}$), it was inaccurate and unstable for Trp-cage (Figure S19). While we also attempted simulations with larger models, we did not search for an optimal $\lambda$ for these. Note that we also did not search for an optimal $\lambda$ when pre-training SchNet to fit $E_{\mathrm{GBn2}}$, as we did not run ML-MD simulations with pre-trained models. We instead simply used our initially selected value.

Table S7: Hyperparameters explored for SchNet.[S22] Bolded values indicate hyperparameters for the best performing model as determined from re-weighted free energy profiles (see Figure 5), while underlined values indicate the hyperparameters of the smaller model used in ML-MD simulation.

| SchNet hyperparameters | |
|---|---|
| Embedding width | **<u>32</u>**, 40, 48, 64, 128 |
| $r_{\text{cut}}$ (nm) | 1, 1.2, 1.4, 1.6, <u>1.8</u>, 2, 3, 4, **5** |
| $N_{\text{IB}}$ | 2, **<u>3</u>**, 4, 5, 6 |

Table S8: Hyperparameters explored for training set-up. Hyperparameters are indicated for both pre-training (i.e., training SchNet to fit $E_{\text{GBn2}}$) and potential contrasting. Bolded values indicate optimal hyperparameters. In cases where only a single value is listed, no further optimization was necessary beyond the initially selected value.

| Hyperparameter | Pre-training | Potential contrasting |
|---|---|---|
| $\zeta$ | $\mathbf{1 \times 10^{-3}}$ | $\mathbf{1 \times 10^{-4}}$ |
| $\mathbf{\Delta F}_k$ $\zeta^a$ | - | **0.75** |
| $\zeta$ decay | **×0.1 / 15 epochs**$^b$ | **×0.1 / 10 epochs** |
| # epochs | **120** | **30** |
| $\lambda$ ($\times 10^{-7}$) | **50** | 0.1, 1, **10**, 50, 100, 500, 1,000, 10,000 |
| Batch size | 10, **20**, 25, 50 | **10**, 20, 25 |
| Frozen parameters | **none** | none, **interaction blocks** |
| Testing set size$^c$ | **50,000** | - |
| Activation | **shifted softplus**[S22] | |
| Optimizer | **Adam**[S23] | |

$^a$ Separate learning rate used for $\mathbf{\Delta F}_k$'s.
$^b$ After the first 60 epochs, pre-train $\zeta$ was reset to $1 \times 10^{-3}$. The same decay scheme was used for the final 60 epochs.
$^c$ Total # of configurations held out of the training set for each protein. Different random sets of configurations were held out every epoch.

#### Software versions & hardware

All models were trained using PyTorch[S19] ver. 1.10.2 and PyTorch Geometric[S20] ver. 2.0.4. This version of PyG contains PyTorch Cluster ver. 1.6.0 and PyTorch Scatter ver. 2.0.9. Both pre-training and potential contrasting were performed across two Nvidia Volta V100 GPUs using the `DistributedDataParallel` class. All configurations for three proteins were loaded onto each individual GPU.

### Noise potential & free energy profile calculations

The multi-state Bennett acceptance ratio[S24] (MBAR) and the unbinned weighted histogram analysis method[S25] (UWHAM), as implemented in the FastMBAR package,[S26] were respectively used to compute the noise potential $u_q(\boldsymbol{x})$ as defined in the potential contrasting loss function (Eq. S3) and the free energy profile from the states sampled in biased umbrella sampling simulations. Free energy profiles were also computed from long timescale unbiased MD simulations. The methods used to compute $u_q(\boldsymbol{x})$ and these free energy profiles are detailed in the subsections below.

#### Constructing the noise potential using FastMBAR

As detailed in section *Umbrella sampling simulation set-up* and shown in equation S1, each window in RMSD-biased umbrella sampling simulations has a different biasing potential $u^i_{\text{bias}}(\boldsymbol{x})$ applied. The potential energy of the $i$th umbrella window in simulations with the GBn2 implicit solvent model is

$$u_i(\boldsymbol{x}) = U_{\text{ff}}(\boldsymbol{x}) + E_{\text{GBn2}}(\boldsymbol{x}) + u^i_{\text{bias}}(\boldsymbol{x}), \tag{S6}$$

where $U_{\text{ff}}(\boldsymbol{x})$ is the vacuum potential energy of the solute computed from the chosen protein force field and $E_{\text{GBn2}}(\boldsymbol{x})$ is the solvation free energy computed from GBn2.

Since a different potential energy function $u_i(\boldsymbol{x})$ was used for each umbrella window, a different distribution was sampled in each window. However, for training with potential contrasting,[S10] the noise samples must be drawn from a single noise distribution $q(\boldsymbol{x})$ yielded by a noise potential $u_q(\boldsymbol{x})$. As explained in the works of Ding et al.[S26] and Ding and Zhang[S10], by pooling all configurations from each umbrella window, they can treated as if they had been drawn from a single generalized ensemble $p_{\text{gm}}(\lambda = i, \boldsymbol{x}) \propto \exp\left(-\beta\left[u_i(\boldsymbol{x}) + v_i\right]\right)$. Here, $\lambda$ is simply a discrete value representing the index of the $i$th state of $p_{\text{gm}}(\lambda = i, \boldsymbol{x})$. The marginal probability density $p_{\text{gm}}(\boldsymbol{x})$ on $\boldsymbol{x}$ in $p_{\text{gm}}(\lambda = i, \boldsymbol{x})$ is yielded by the potential energy

$$u_{\text{gm}}(\boldsymbol{x}) = -\beta^{-1} \ln \sum_{i=1}^{M} \exp\left(-\beta\left[u_i(\boldsymbol{x}) + v_i\right]\right), \tag{S7}$$

where $v_i$ is a fitted biasing energy and $M$ is the total number of states. Fitting of $v_i$ is conducted so that the relative free energy $G_i$ of the $i$th state of $p_{\text{gm}}(\lambda = i, \boldsymbol{x})$ matches the relative population of configurations in the $i$th state of $p_{\text{gm}}(\lambda = i, \boldsymbol{x})$. Thus, $v_i$ must satisfy the condition $G_i = G_i^* + v_i = -\ln\left[n_i/N_{\text{tot}}\right]$, where $G_i^*$ is the relative free energy of the protein-solvent system in the $i$th thermodynamic state / umbrella window with the potential $u_i(\boldsymbol{x})$. These $v_i$'s are fit by minimizing the convex objective function

$$f(v_1, v_2, ..., v_M) = \frac{1}{N_{\text{tot}}} \sum_{j=1}^{M} \sum_{k=1}^{n_j} \ln\left[\sum_{l=1}^{M} \exp\left(-\beta\left[u_l\left(\boldsymbol{x}_j^k\right) + v_l\right]\right)\right] + \sum_{i=1}^{M} \frac{n_i}{N_{\text{tot}}} v_i, \tag{S8}$$

where here $N_{\text{tot}} = \sum_{i=1}^{M} n_i$. By solving for $v_i$ we can compute $u_{\text{gm}}(\boldsymbol{x})$ as shown in equation S7 and treat the marginal probability density $p_{\text{gm}}(\boldsymbol{x})$ as our noise distribution $q(\boldsymbol{x})$. Thus, $u_{\text{gm}}(\boldsymbol{x})$ is our noise potential energy $u_q(\boldsymbol{x})$.

In practice, fitting $v_i$ and computing $u_q(\boldsymbol{x})$ was done using the FastMBAR package.[S26] This process of fitting $v_i$ to compute $u_q(\boldsymbol{x})$ was done for each protein in our training set. As such, the $k$th protein has its own noise distribution $q_k(\boldsymbol{x})$ yielded by $u_q^k(\boldsymbol{x})$. These $u_q^k(\boldsymbol{x})$'s were used in the potential contrasting objective function (Eq. S3).

#### Free energy profile calculations

Free energy profiles were computed with respect to backbone heavy atom RMSD from the folded configuration $\boldsymbol{x}_\circ$ and with respect to the radius of gyration (Rg) of the backbone heavy atoms. The methods used to compute these free energy profiles are detailed below.

#### Calculating free energy profiles from umbrella simulations using FastMBAR

Computing $v_i$ and $u_q(\boldsymbol{x})$ as detailed in section *Constructing the noise potential using FastMBAR* will enable usage of a simple re-weighting scheme to compute the free energy profile. While the following text details the computation of RMSD free energy profiles, the same procedure is used to compute Rg free energy profiles. The RMSD range can be split into $L$ different bins, with $\text{RMSD}_l$ representing the central RMSD of the $l$th bin. $L$ was selected so that the width of the $l$th bin was 0.05 nm RMSD (or 0.0125 nm Rg). An energy of 0 or $+\infty$ kJ/mol is assigned to each configuration $\boldsymbol{x}$ according to the equation

$$R_l(\boldsymbol{x}) = \begin{cases} 0, & \text{if } \text{RMSD}(\boldsymbol{x}, \boldsymbol{x}_\circ) \in [\text{RMSD}_{l-0.5}, \text{RMSD}_{l+0.5}] \\ +\infty, & \text{otherwise.} \end{cases}, \tag{S9}$$

where $\text{RMSD}_{l-0.5}$ and $\text{RMSD}_{l+0.5}$ are the lower and upper edges of the $l$th RMSD bin, respectively. After computing $R_l(\boldsymbol{x})$, we can compute the relative free energy of the $l$th bin

$$F_l = -\beta^{-1} \ln \left[ \frac{1}{N_{\text{tot}}} \sum_{j=1}^{M} \sum_{k=1}^{n_j} \exp\left(-\beta \left[ R_l\left(\boldsymbol{x}_j^k\right) - u_q\left(\boldsymbol{x}_j^k\right) \right]\right) \right], \tag{S10}$$

where the summation runs over all configurations from all $M$ umbrella windows.

For all free energy profiles computed in this way, the standard deviations were computed using block bootstrapping.[S27] Based on the save frequency of each simulation, the block size was selected as the number of consecutive configurations representing 2 ns of simulation. Ten bootstrap samples were constructed to compute each free energy profile.

#### Calculating free energy profiles from unbiased explicit solvent simulations

For protein configurations generated from long timescale, unbiased explicit solvent MD simulations, we computed free energy profiles directly from RMSD or Rg histograms with $L$ total bins. $L$ was selected so that the width of the $l$th bin was 0.05 nm RMSD or 0.0125 nm

Rg. This can be done by first computing the number of configurations $n_l$ that fall within the RMSD or Rg range of the $l$th bin, next estimating the probability of configurations occurring within the $l$th bin as $p_l = n_l/N_{\text{tot}}$, and lastly by computing the relative free energy of the $l$th bin as $F_l = -\ln p_l$.

#### TICA free energy profiles

Following the procedure of Ojaghlou et al.[S12], dimensionality reduction was performed using time-lagged independent component analysis (TICA) on the explicit solvent configurations. Each backbone dihedral angle $\omega$ was featurized as pairs of $\sin(\omega)$ and $\cos(\omega)$, a lag time of 10 ns was selected, and the kinetic map option was set to false. The two outputted time-lagged independent components (TICs) were then binned to evenly spaced points on a grid matrix $L = (20 \times 20)$, and a kernel density estimate was computed over the grid points. At each individual grid point $\boldsymbol{X}_{i,j} = (x_i, y_j)$, the probability density

$$p_{i,j}\left(\boldsymbol{X}_{i,j}\right) = \frac{1}{N_c} \sum_{l=1}^{L_y} \sum_{k=1}^{L_x} c_{k,l} K_{\boldsymbol{H}}\left(\boldsymbol{X}_{i,j}, \boldsymbol{X}_{k,l}\right) \tag{S11}$$

was evaluated. Here, $c_{k,l}$ is simply the sum of the weights of each configuration assigned to the bin centered at $\boldsymbol{X}_{k,l}$, $N_c$ is the sum of the weights of all configurations, and $K_{\boldsymbol{H}}\left(\boldsymbol{X}_{i,j}, \boldsymbol{X}_{k,l}\right)$ is a bivariate Gaussian kernel function with the bandwidth matrix $\boldsymbol{H}$ computed using Scott's Rule.[S28,S29] All explicit solvent configurations were uniformly weighted. The free energy for the bin centered at $\boldsymbol{X}_{i,j}$ was computed as $F_{i,j} = -\ln p_{i,j}$.

Free energy profiles were created for the GBn2 and SchNet implicit solvent umbrella sampling configurations by first fitting the backbone dihedrals from these configurations to the TICs computed for the explicit solvent data. Weights were then computed for each configuration. Following the procedure used to generate the noise potential $u_q(\boldsymbol{x})$ detailed in section *Constructing the noise potential using FastMBAR*, a generalized ensemble $p_{\text{gm}}(\lambda = i, \boldsymbol{x}) \propto \exp\left(-\beta\left[u_i(\boldsymbol{x}) + v_i\right]\right)$ was constructed and the energy of the corresponding marginal probability density $p_{\mathrm{gm}}(\boldsymbol{x})$ on $\boldsymbol{x}$ in $p_{\mathrm{gm}}(\lambda = i, \boldsymbol{x})$ was computed (Eq. S7). We then computed a weight $w(\boldsymbol{x})$ for each configuration. For configurations from GBn2 implicit solvent, $w(\boldsymbol{x}) = \frac{p(\boldsymbol{x})}{p_{\mathrm{gm}}(\boldsymbol{x})} = \exp\left(-\beta\left[U_{\mathrm{C36}}(\boldsymbol{x}) + E_{\mathrm{GBn2}}(\boldsymbol{x}) - u_{\mathrm{gm}}(\boldsymbol{x})\right]\right)$. For configurations from the SchNet implicit solvent $w(\boldsymbol{x}) = \frac{p(\boldsymbol{x};\boldsymbol{\theta}^*)}{p_{\mathrm{gm}}(\boldsymbol{x};\boldsymbol{\theta}^*)} = \exp\left(-\beta\left[U_{\mathrm{C36}}(\boldsymbol{x}) + E_{\mathrm{SchNet}}(\boldsymbol{x};\boldsymbol{\theta}^*) - u_{\mathrm{gm}}(\boldsymbol{x};\boldsymbol{\theta}^*)\right]\right)$. Using these weights, free energy profiles were created by following the same procedure used to generate TICA free energy profiles for the JIP1 10-mer explicit solvent configurations.

### Figures

All figures referenced in the main text (Figures S1 to S11) and all figures referenced exclusively in this Supporting Information document (Figures S12 to S19) are displayed below.

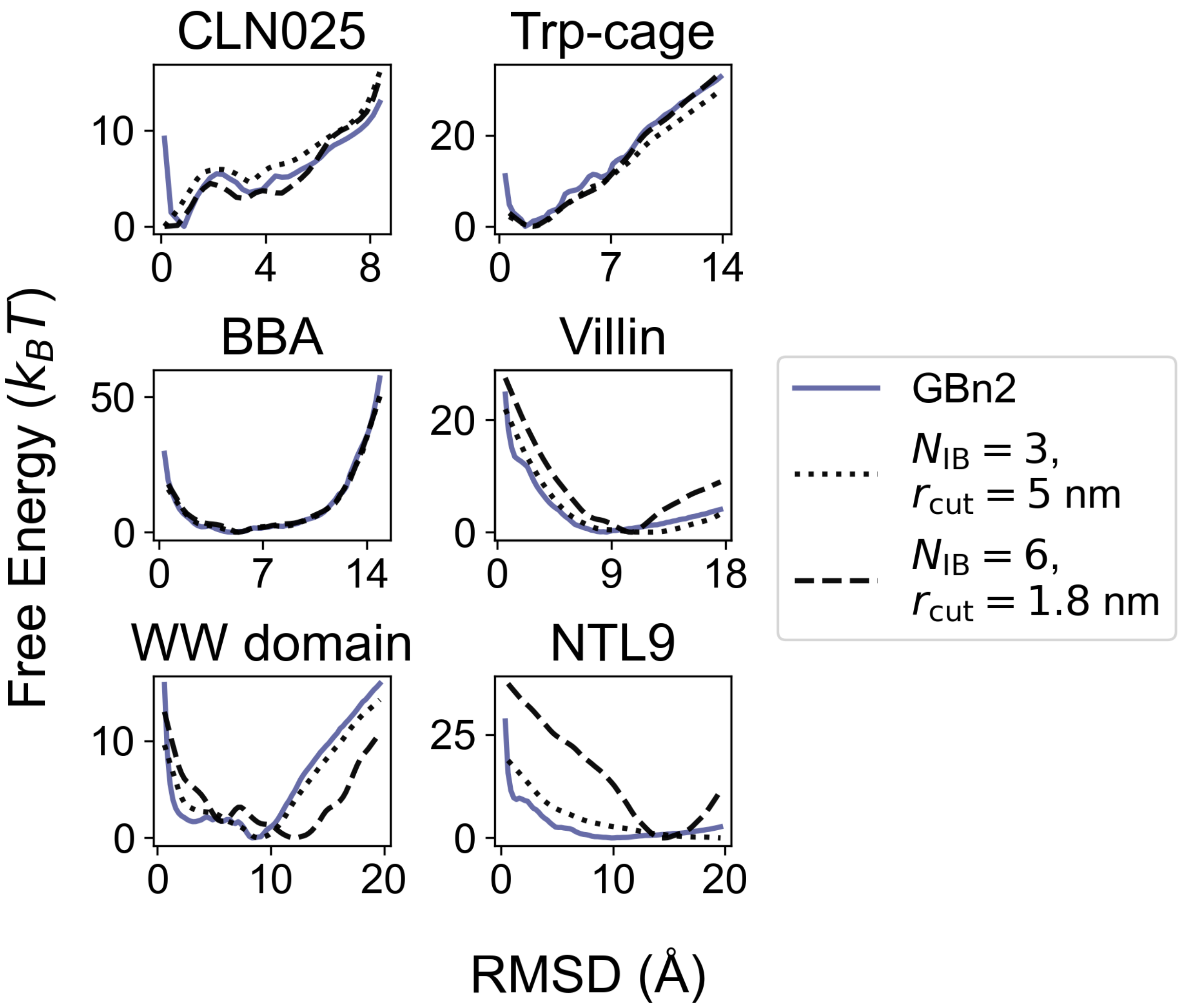


Figure S1: A SchNet model[S22] with larger $r_{\mathrm{cut}}$ but smaller $N_{\mathrm{IB}}$ outperforms a model with smaller $r_{\mathrm{cut}}$ but larger $N_{\mathrm{IB}}$. Both models were trained to fit $E_{\mathrm{GBn2}}$. Free energy profiles for the model with larger $r_{\mathrm{cut}}$ are shown in the dotted line, while free energy profiles for the model with a larger $N_{\mathrm{IB}}$ are shown in the dashed lines. For reference, the free energy profiles computed from umbrella simulations with the GBn2 implicit solvent model are also shown as solid lines. The $x$ axes in the plots correspond to the RMSD from the folded structures. Results for the SchNet model were determined using a re-weighting scheme as detailed in the *Methods* Section.

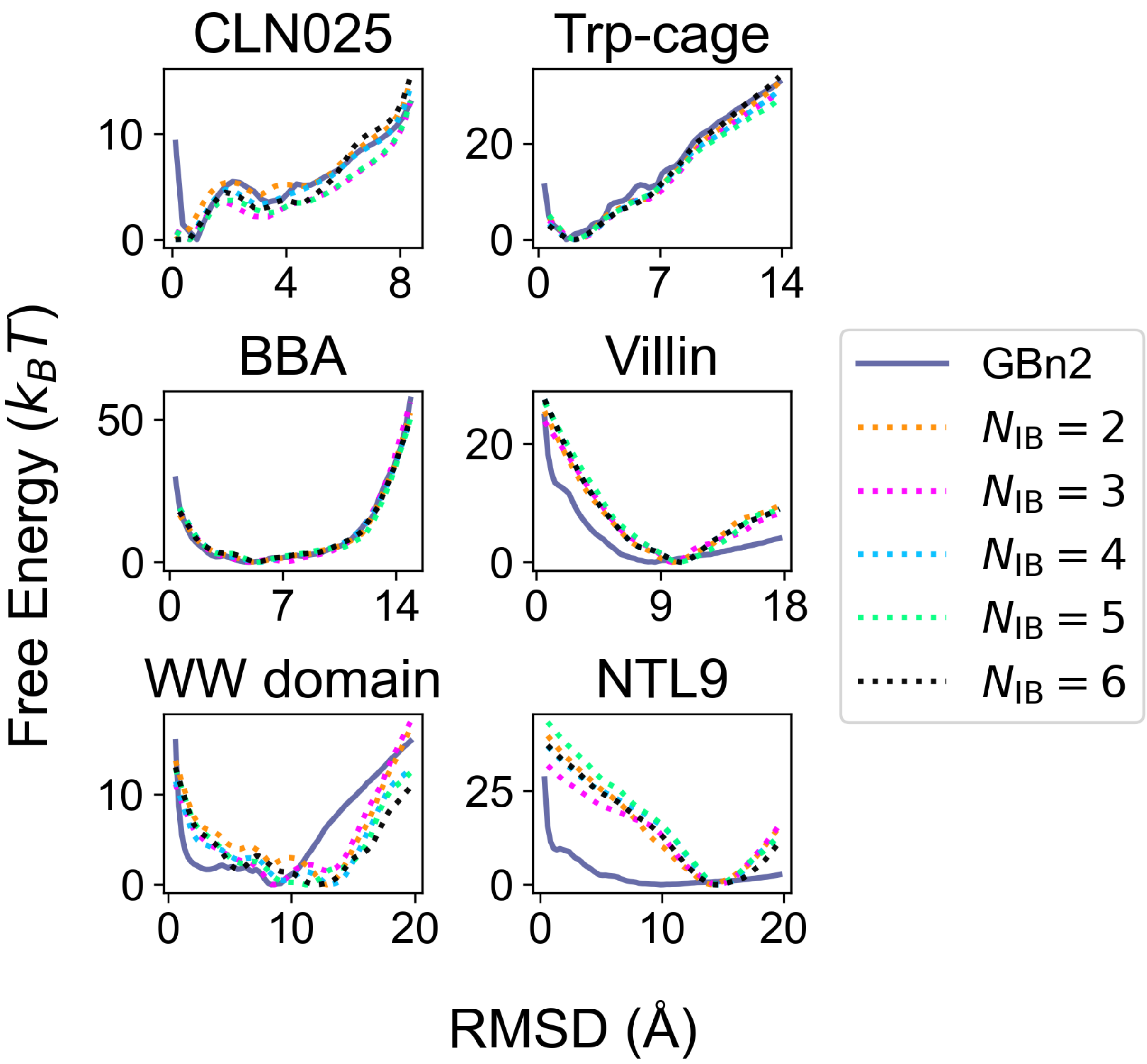


Figure S2: SchNet models trained to fit $E_{\text{GBn2}}$ with different values of $N_{\text{IB}}$ perform similarly. Free energy profiles are shown for all models. The cutoff distance is held constant at $r_{\text{cut}} = 1.8$ nm. The $x$ axes in the plots correspond to the RMSD from the folded structures. For reference, the free energy profiles computed from umbrella simulations with the GBn2 implicit solvent model are also shown as solid lines. Results for the SchNet model were determined using a re-weighting scheme as detailed in the *Methods* Section.

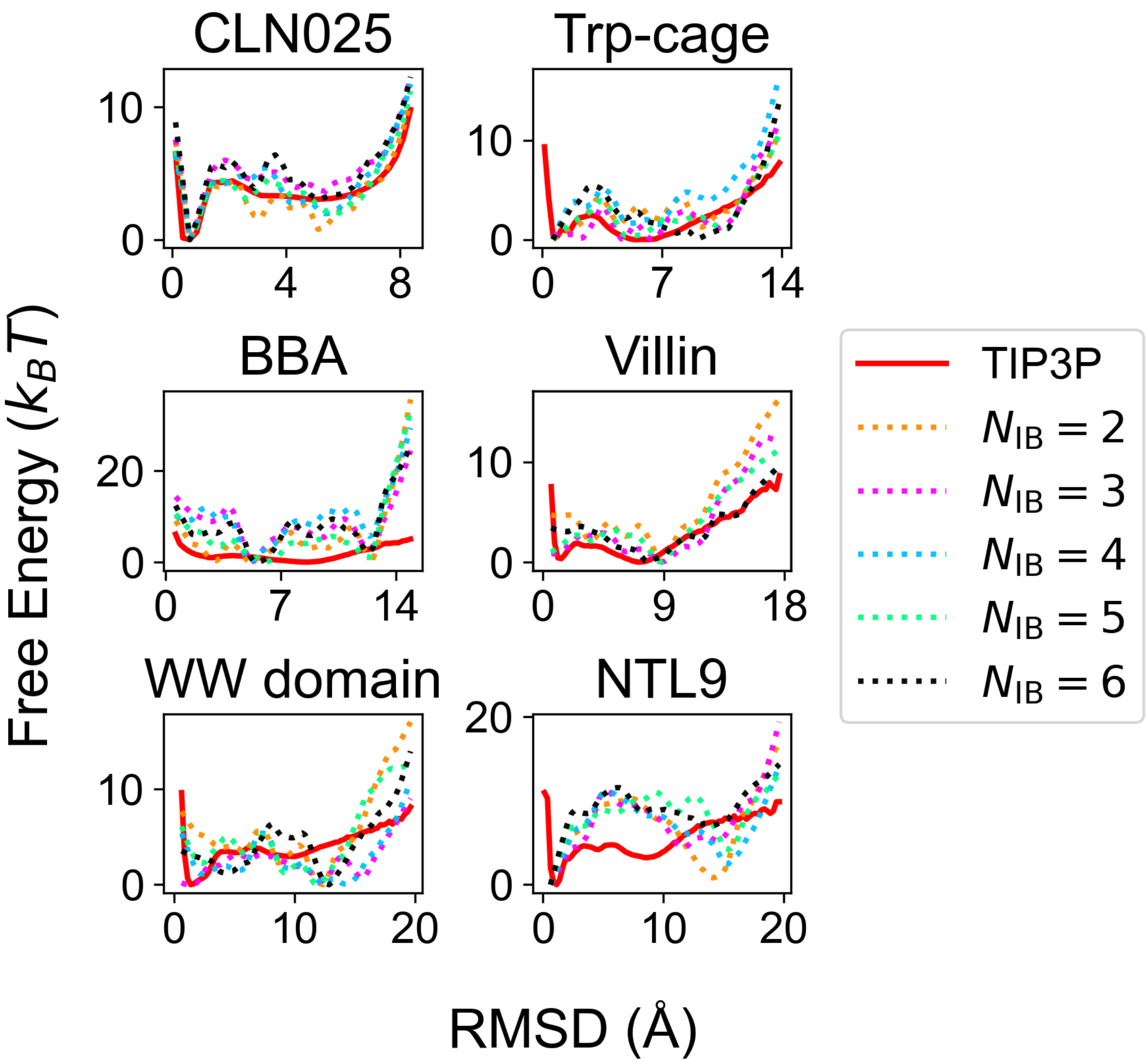


Figure S3: Free energy profiles for SchNet models trained with potential contrasting with different values of $N_{\mathrm{IB}}$. The cutoff distance is held constant at $r_{\mathrm{cut}} = 1.8$ nm. The $x$ axes in the plots correspond to the RMSD from the folded structures. For reference, the free energy profiles computed from simulations conducted by Lindorff-Larsen et al.[S2] in TIP3P explicit solvent are shown as red solid lines. Results for the SchNet model were determined using a re-weighting scheme as detailed in the *Methods* Section.

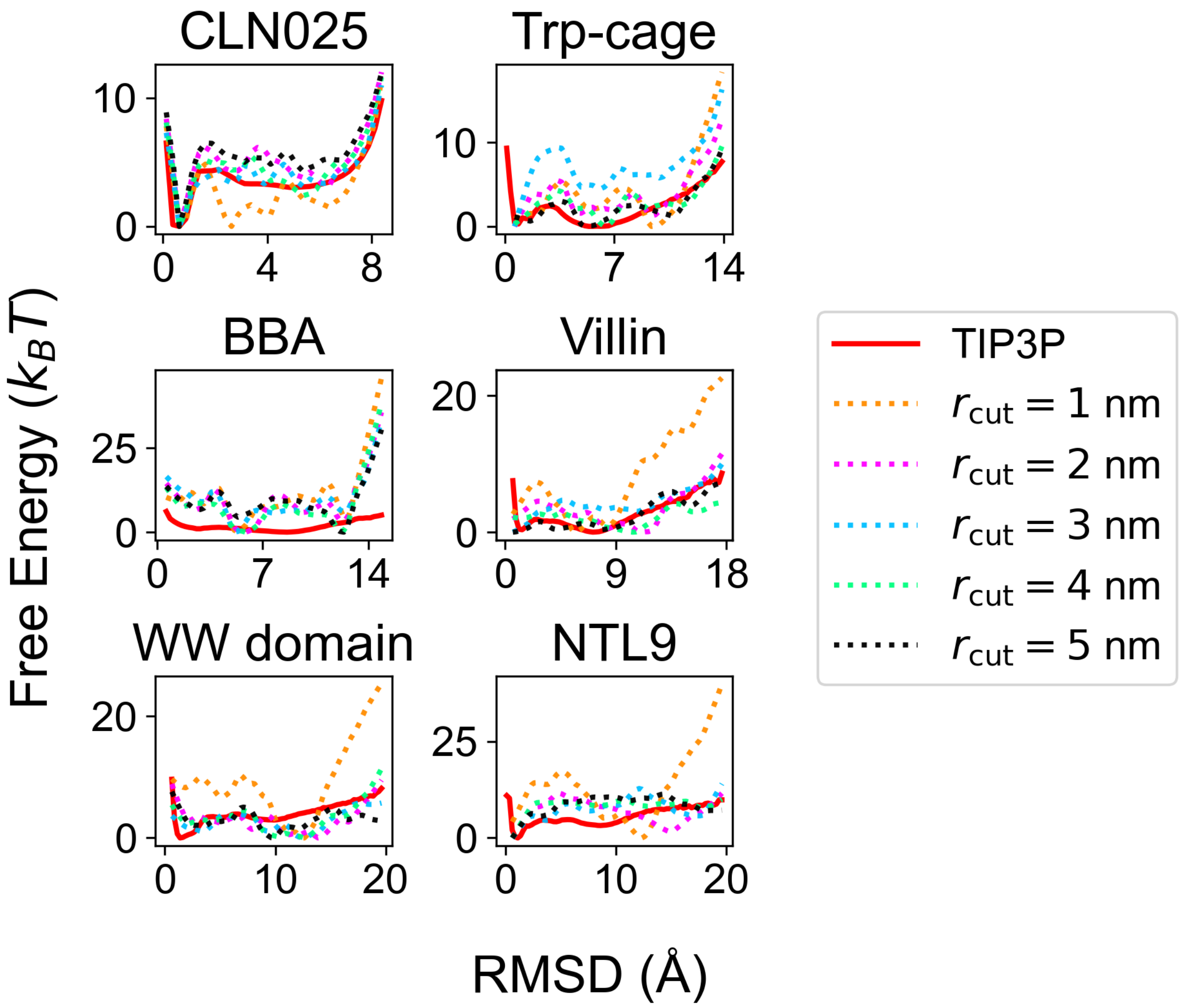


Figure S4: Free energy profiles for SchNet models trained with potential contrasting with different values of $r_{\mathrm{cut}}$. The number of interaction blocks is held constant at $N_{\mathrm{IB}} = 3$. The $x$ axes in the plots correspond to the RMSD from the folded structures. For reference, the free energy profiles computed from simulations conducted by Lindorff-Larsen et al.[S2] in TIP3P explicit solvent are shown as red solid lines. Results for the SchNet models were determined using a re-weighting scheme as detailed in the *Methods* Section.

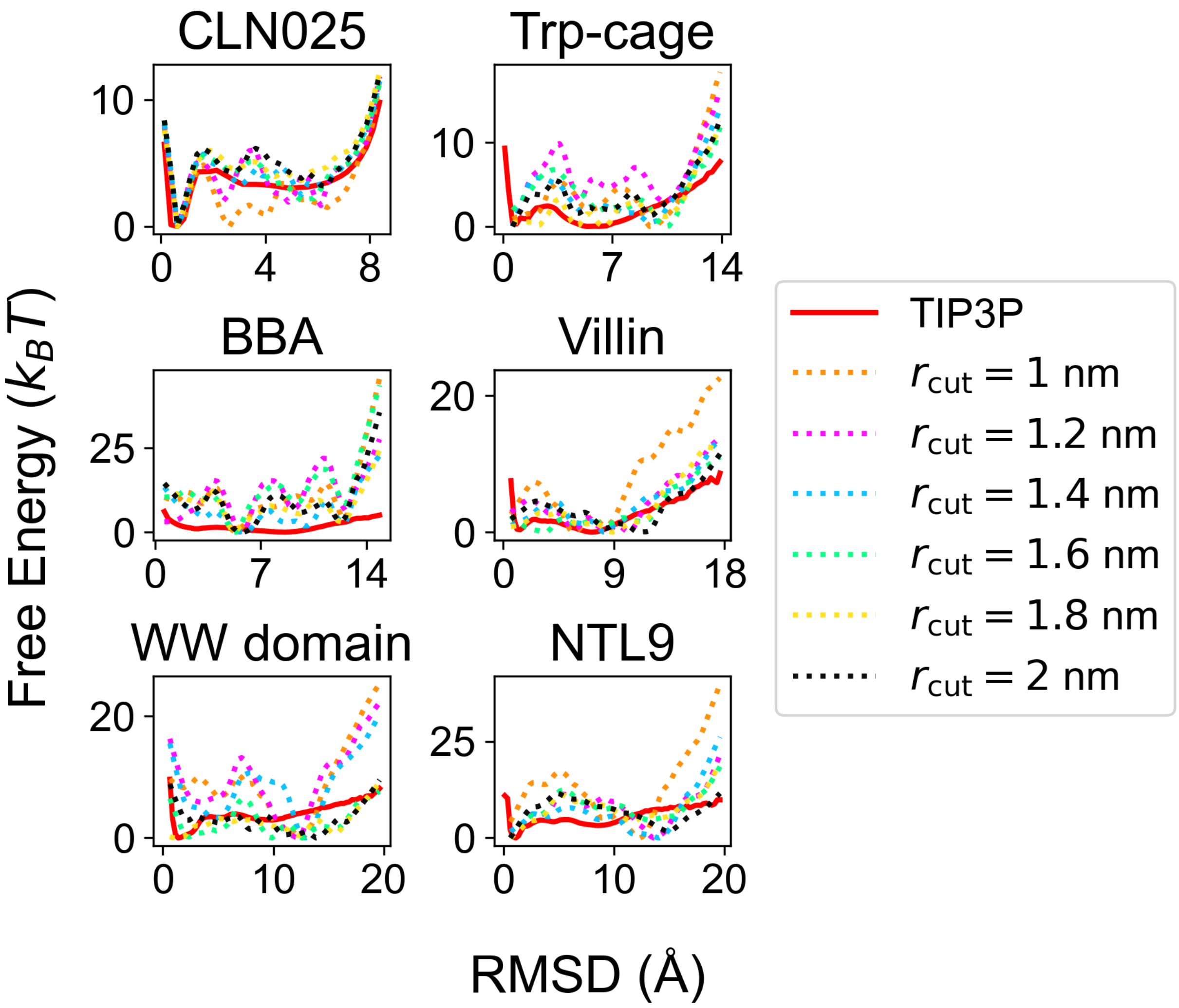


Figure S5: Free energy profiles for SchNet models trained with potential contrasting with values of $r_{\text{cut}}$ from 1 to 2 nm. The number of interaction blocks is held constant at $N_{\text{IB}} = 3$. The $x$ axes in the plots correspond to the RMSD from the folded structures. For reference, the free energy profiles computed from simulations conducted by Lindorff-Larsen et al.[S2] in TIP3P explicit solvent are shown as red solid lines. Results for the SchNet models were determined using a re-weighting scheme as detailed in the *Methods* Section.

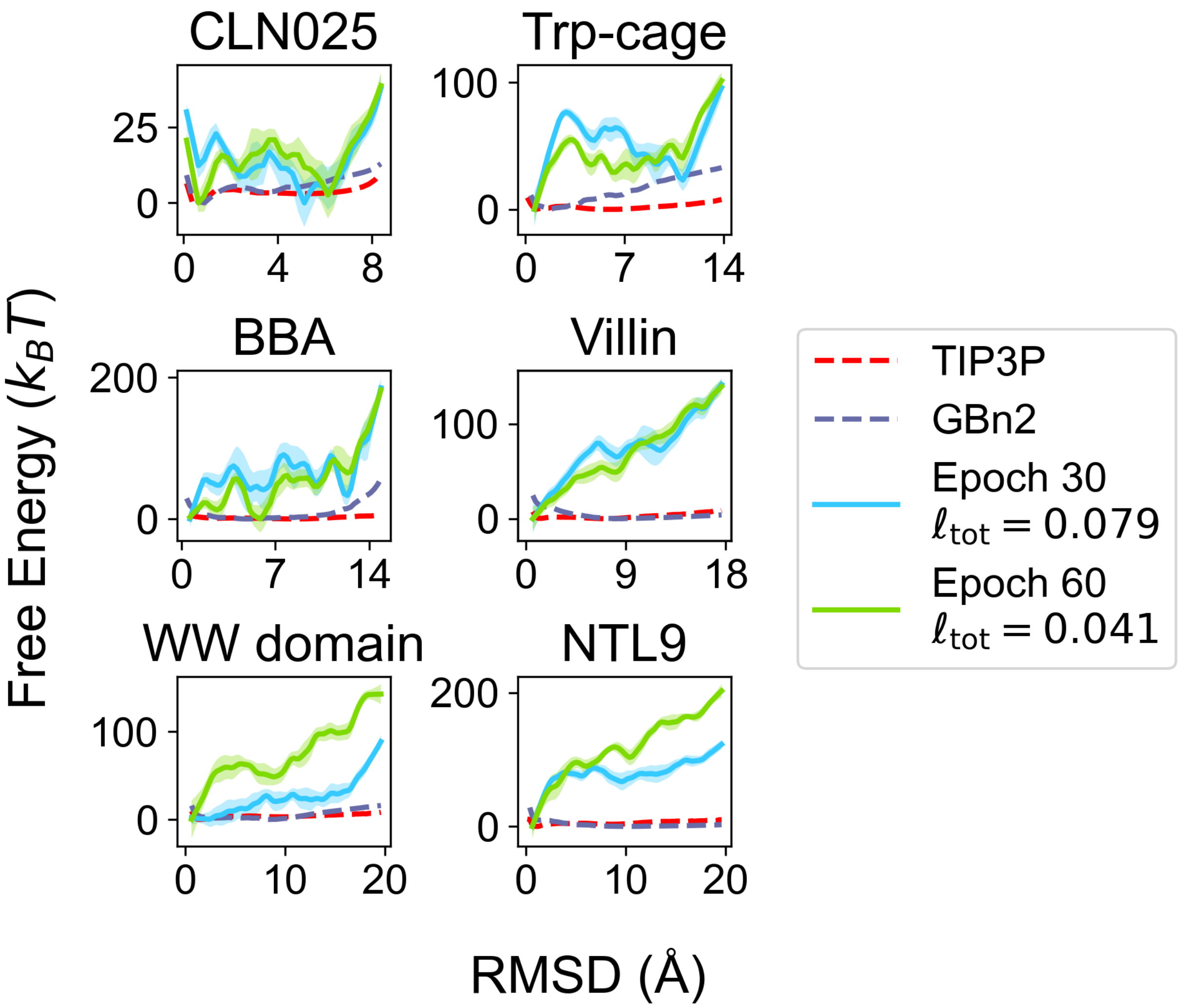


Figure S6: A SchNet model ($r_{\mathrm{cut}} = 1.8$ nm, $N_{\mathrm{IB}} = 3$) trained with the potential contrasting method without prior pre-training fails to match the free energy profiles from explicit solvent simulations. Free energy profiles for the model trained for 30 and 60 epochs are shown in light blue and light green, respectively. The $x$ axes in the plots correspond to the RMSD from the folded structures. Performance is still poor despite the model trained for 60 epochs reaching a similar value of the potential contrasting loss function ($\ell_{\mathrm{tot}} = 0.041$) to the model used for ML-MD simulations in the main text ($r_{\mathrm{cut}} = 1.8$ nm, $N_{\mathrm{IB}} = 3$; $\ell_{\mathrm{tot}} = 0.038$). For reference, the free energy profiles computed from simulations conducted by Lindorff-Larsen et al.[S2] in TIP3P explicit solvent are shown as red dashed lines, and the free energy profiles computed from umbrella simulations with the GBn2 implicit solvent model are shown as blue dashed lines. Results for the SchNet model were determined using a re-weighting scheme as detailed in the *Methods* Section.

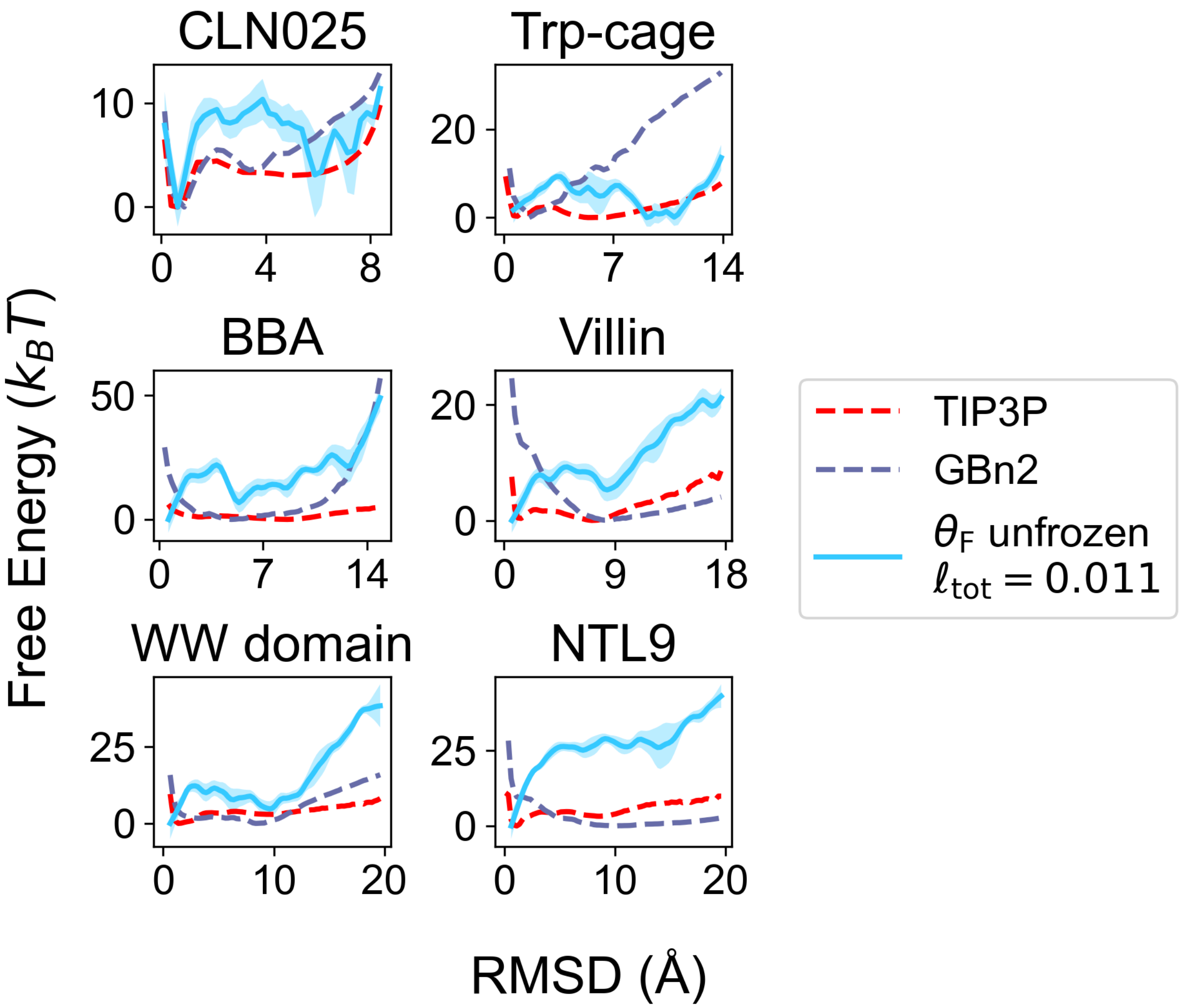


Figure S7: A SchNet model ($r_{cut} = 1.8$ nm, $N_{IB} = 3$) trained with the potential contrasting and pre-training, but without holding the interaction block parameters ($\boldsymbol{\theta_F}$) constant, fails to match the free energy profiles from explicit solvent simulations. The free energy profiles for this model are shown in light blue. The $x$ axes in the plots correspond to the RMSD from the folded structures. Notably, this model reaches a lower value of the potential contrasting loss function ($\ell_{tot} = 0.11$) than the model used for ML-MD simulations in main text ($r_{cut} = 1.8$ nm, $N_{IB} = 3$; $\ell_{tot} = 0.038$). For reference, the free energy profiles computed from simulations conducted by Lindorff-Larsen et al.[S2] in TIP3P explicit solvent are shown as red dashed lines, and the free energy profiles computed from umbrella simulations with the GBn2 implicit solvent model are shown as blue dashed lines. Results for the SchNet model were determined using a re-weighting scheme as detailed in the *Methods* Section.

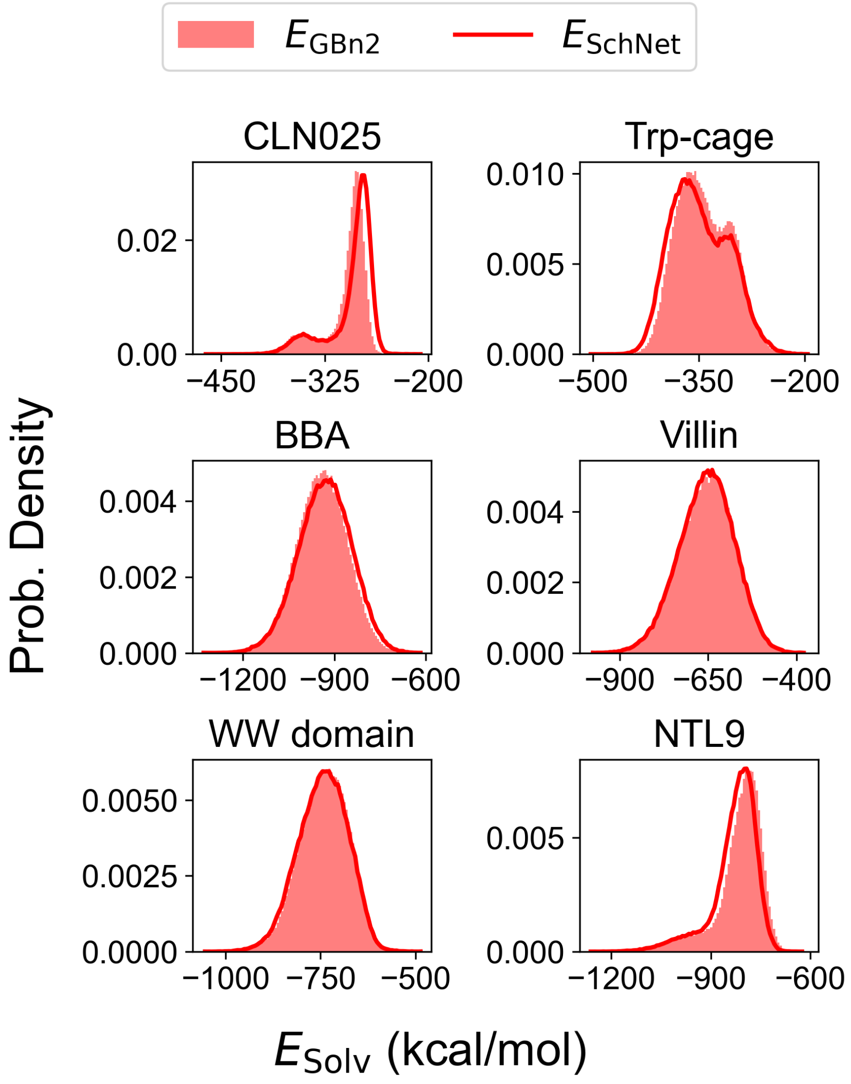


Figure S8: The optimal SchNet model predicts solvation free energy ($E_{\text{SchNet}}$) distributions that are similar in scale to the GBn2 solvation free energy distributions ($E_{\text{GBn2}}$). Solvation free energies are shown for a SchNet model with $r_{\text{cut}} = 1.8$ nm and $N_{\text{IB}} = 3$. The $E_{\text{GBn2}}$ distributions are displayed as filled in, while the $E_{\text{SchNet}}$ distributions are shown as solid lines.

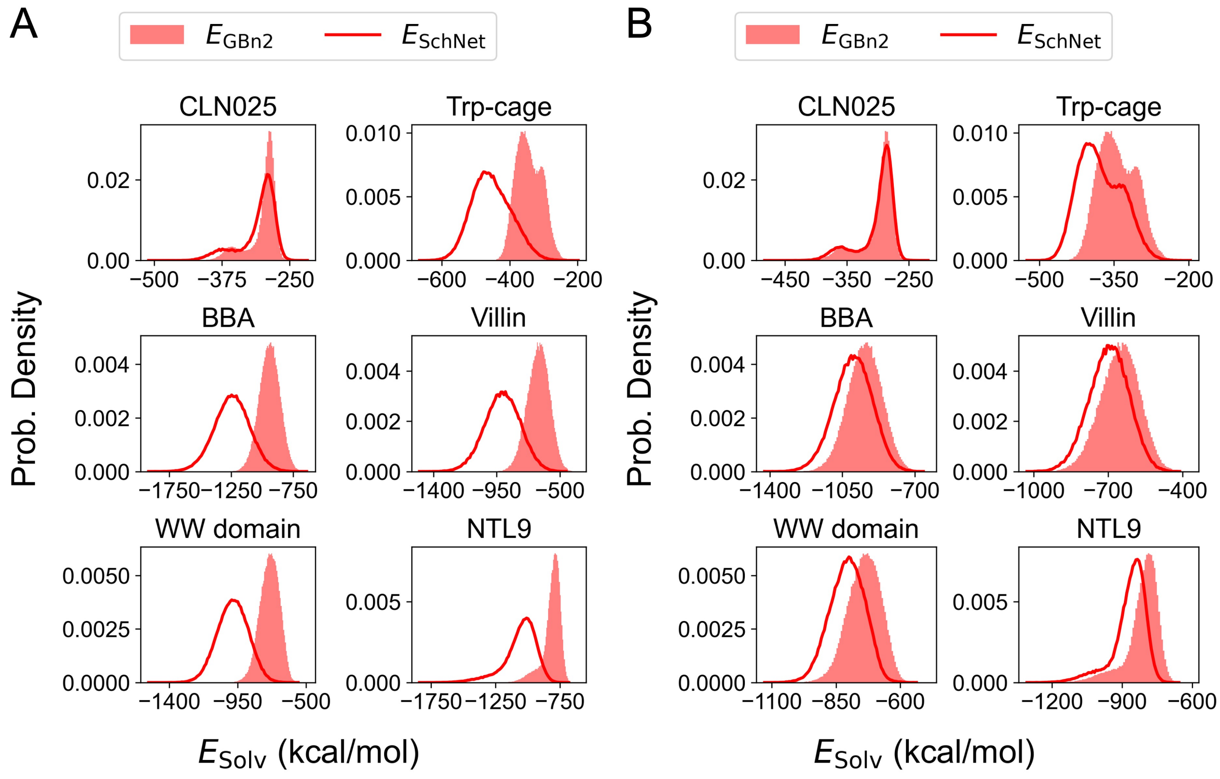


Figure S9: A SchNet implicit solvent trained without pre-training or without holding the interaction block constant yields unrealistic solvation free energies ($E_{\mathrm{SchNet}}$). Solvation free energies are shown for a SchNet model with $r_{\mathrm{cut}} = 1.8$ nm and $N_{\mathrm{IB}} = 3$. The $E_{\mathrm{GBn2}}$ distributions are displayed as filled in, while the $E_{\mathrm{SchNet}}$ distributions are shown as solid lines. (A) $E_{\mathrm{SchNet}}$ distributions for a model that was not pre-trained. For most proteins, SchNet appears to arbitrarily predict low energies. (B) $E_{\mathrm{SchNet}}$ distributions for a model that was pre-trained, but did not have the interaction blocks held constant during training with potential contrasting. Compared to the distributions shown in Figure S8, these distributions are shifted more negatively.

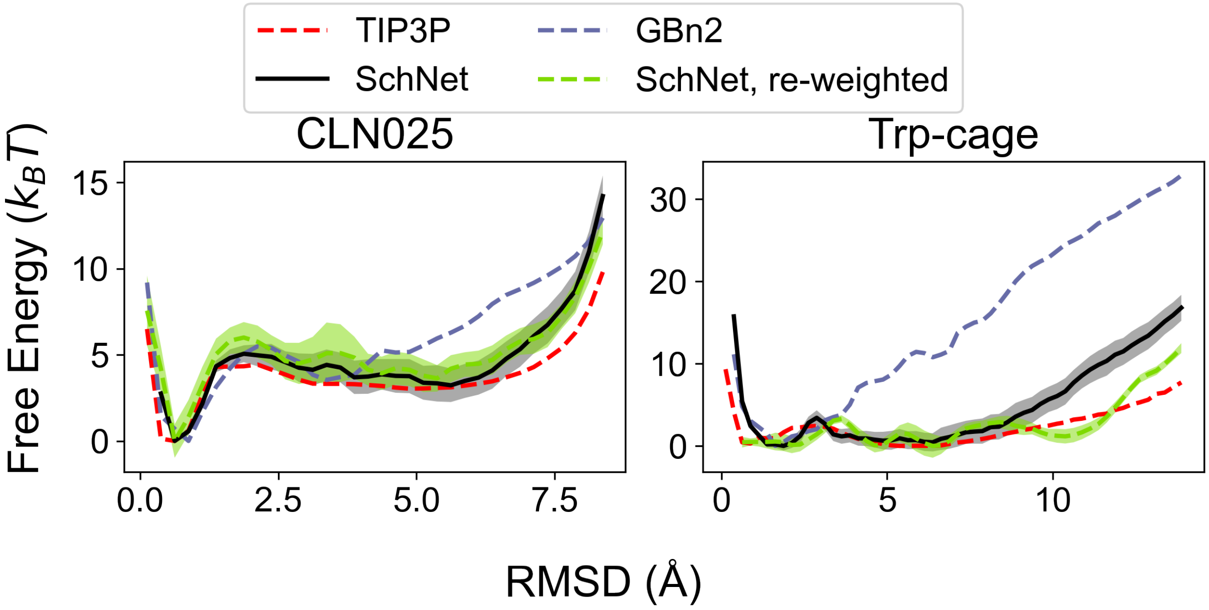


Figure S10: Re-weighted free energy profiles closely match those computed from ML-MD simulations. Explicit solvent (denoted as TIP3P), GBn2, and the SchNet implicit solvent free energy profiles (both re-weighted and from ML-MD simulations) for chignolin CLN025 and Trp-cage are displayed. The $x$ axes in the plots correspond to the RMSD from the folded structures. TIP3P free energy profiles were computed from the configurations generated by Lindorff-Larsen et al.[S2] while GBn2 and SchNet free energy profiles were computed by solving the MBAR equation for configurations from umbrella sampling simulations. The re-weighted SchNet free energy profiles ($r_{\text{cut}}$ = 1.8 nm, $N_{\text{IB}}$ = 3) were computed following the re-weighting procedure detailed in the *Methods* section.

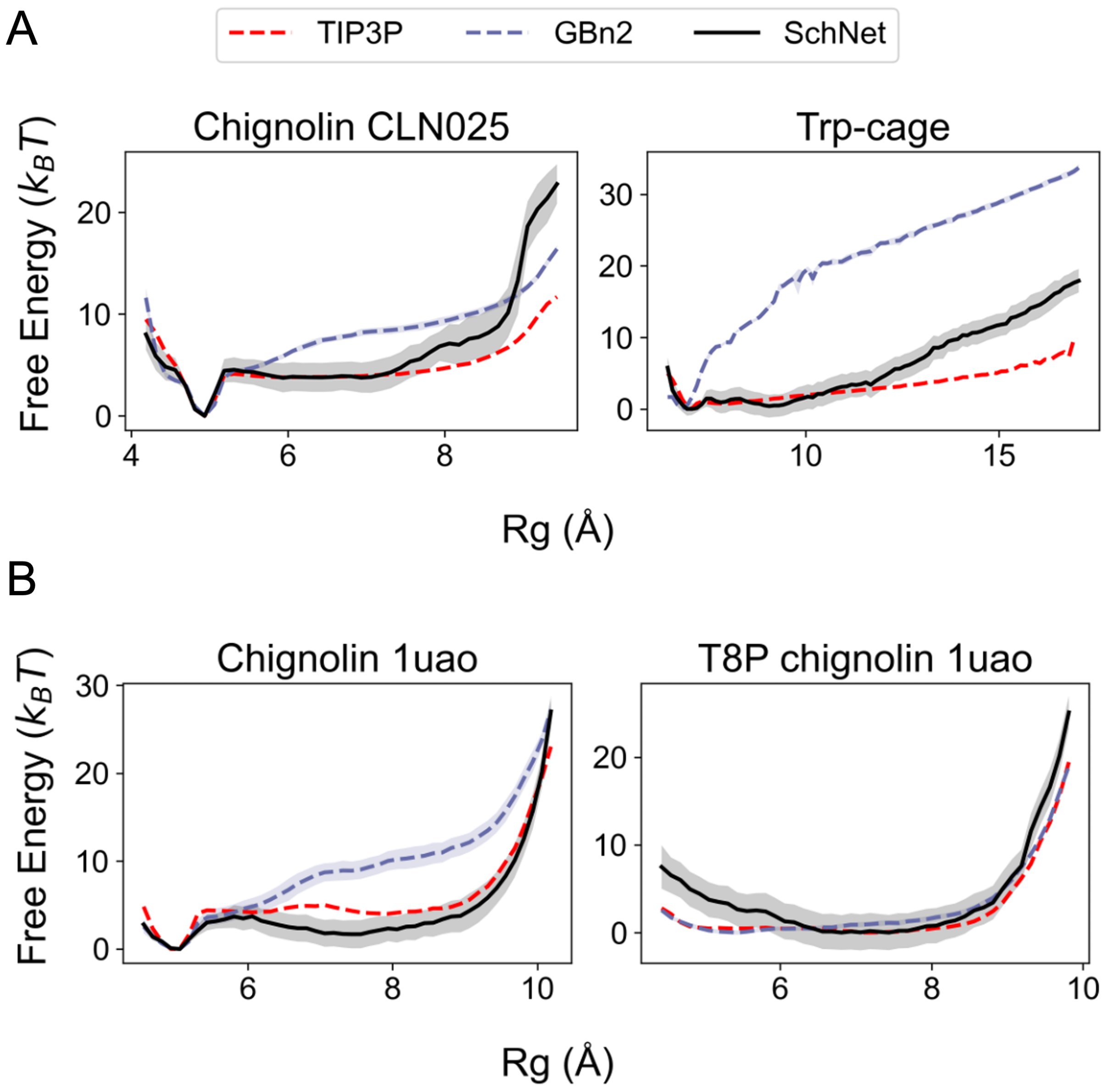


Figure S11: Comparison between the free energy profiles computed from explicit solvent simulations (TIP3P), implicit solvent simulations (GBn2), and the optimized SchNet model. Results for the SchNet model were determined using ML-MD umbrella simulations. The $x$ axes in the plots correspond to the radius of gyration (Rg) of the backbone heavy atoms. (A) Free energy profiles for chignolin CLN025 and Trp-cage. (B) Free energy profiles for chignolin 1uao and a T8P mutant. TIP3P and GBn2 free energy profiles were computed from umbrella simulations.

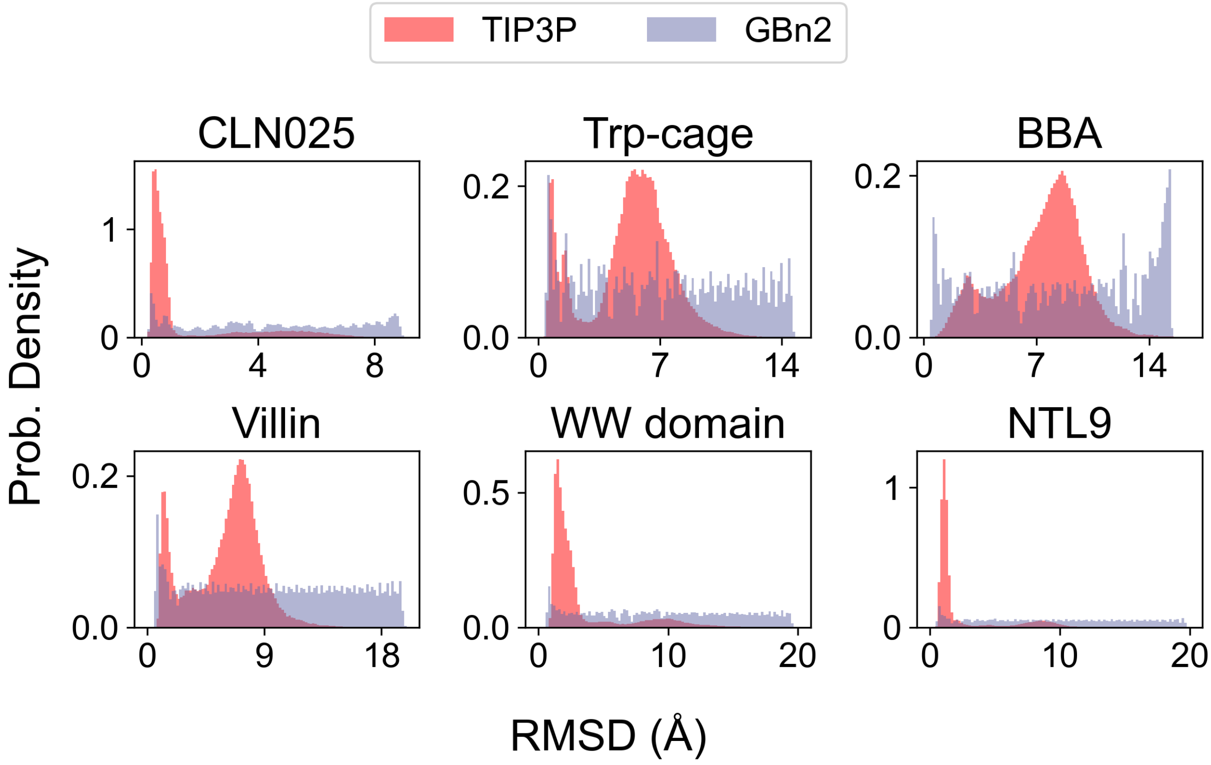


Figure S12: Explicit solvent (denoted as TIP3P) and GBn2 configuration RMSD histograms. RMSD is computed with respect to peptide backbone heavy atom positions from $\boldsymbol{x}_{\circ}$. All TIP3P configurations for these 6 proteins were generated by Lindorff-Larsen et al.[S2] Noise configurations from the GBn2 umbrella sampling ensembles fully cover the RMSD ranges sampled by the TIP3P data configurations, while also including configurations outside of these ranges. This follows best-practice for generating noise samples for training with potential contrasting.[S10]

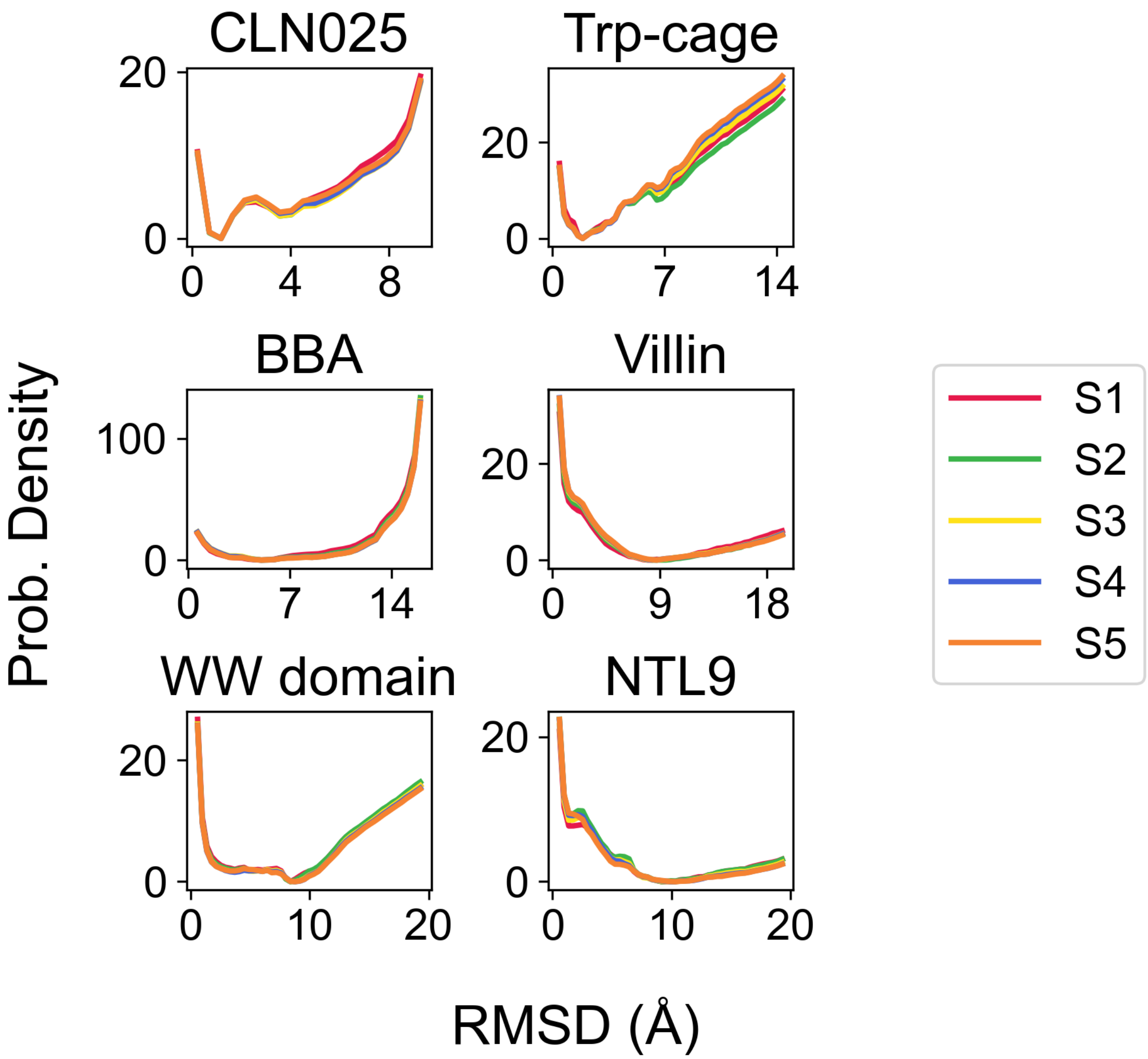


Figure S13: Converged free energy profiles for all training set proteins. Umbrella sampling windows were split 5 ways to show that umbrella sampling GBn2 free energy profiles were converged. These splits were created according the following example for CLN025: S1 0 - 212 ns, S2 0 - 424 ns, S3 0 - 636 ns, S4 0 - 848 ns, and S5 0 - 1,060 ns. For all proteins, the final split includes all configurations for each window.

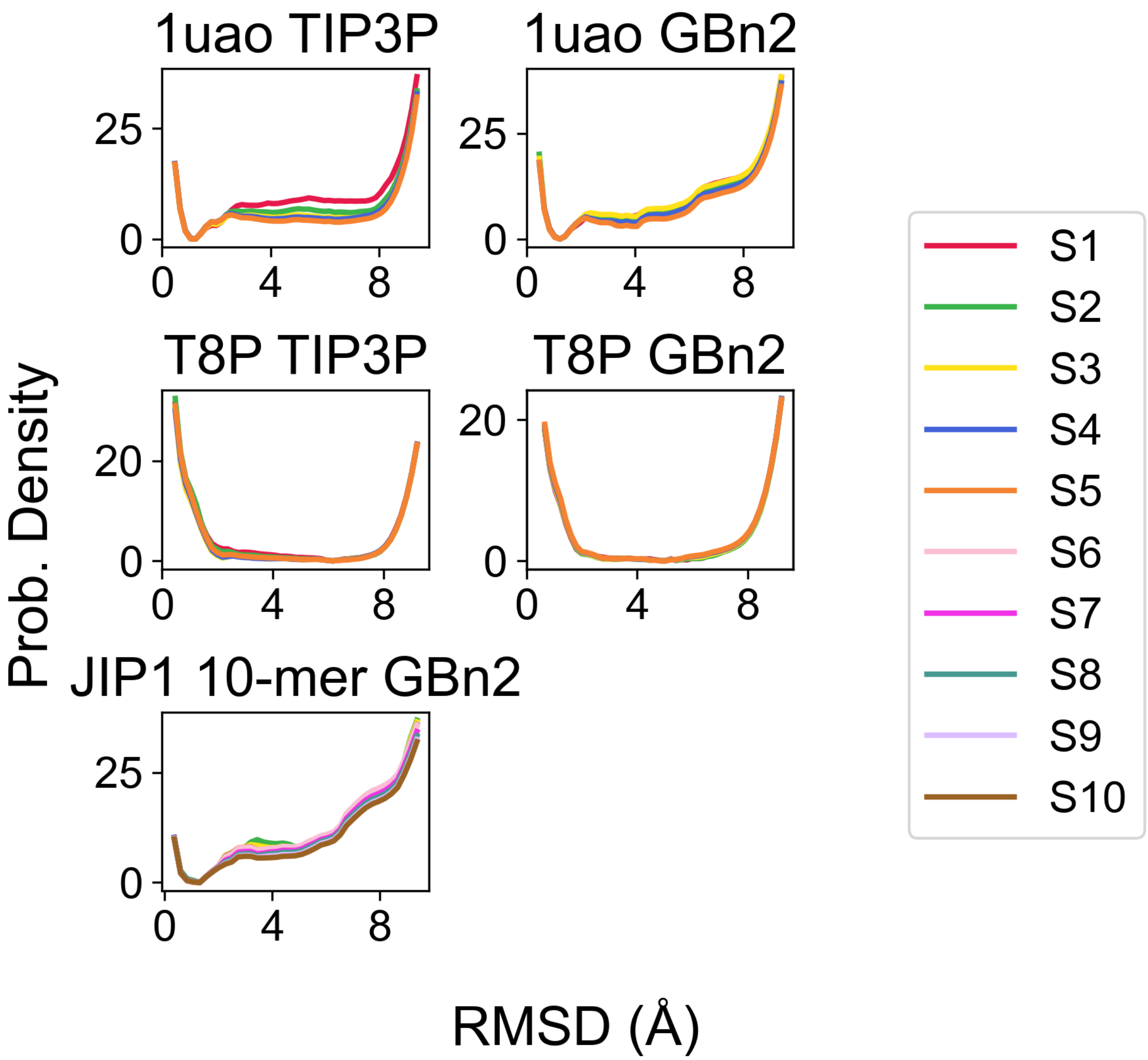


Figure S14: Converged free energy profiles for all testing set proteins. Umbrella sampling windows were split 5 ways for 1uao and T8P and 10 ways for the JIP1 10-mer to show that umbrella sampling TIP3P and GBn2 free energy profiles were converged. For 1uao and T8P, splits were created according the following scheme: S1 0 - 12.5 ns, S2 0 - 25 ns, S3 0 - 37.5 ns, S4 0 - 50 ns, and S5 0 - 62.5 ns. For the JIP1 10-mer, splits were created according to the following scheme: S1 0 - 100 ns, S2 0 - 200 ns, S3 0 - 300 ns, S4 0 - 400 ns, S5 0 - 500 ns, S6 0 - 600 ns, S7 0 - 700 ns, S8 0 - 800 ns, S9 0 - 900 ns, and S10 0 - 1,000 ns. For all proteins, the final split includes all configurations for each window.

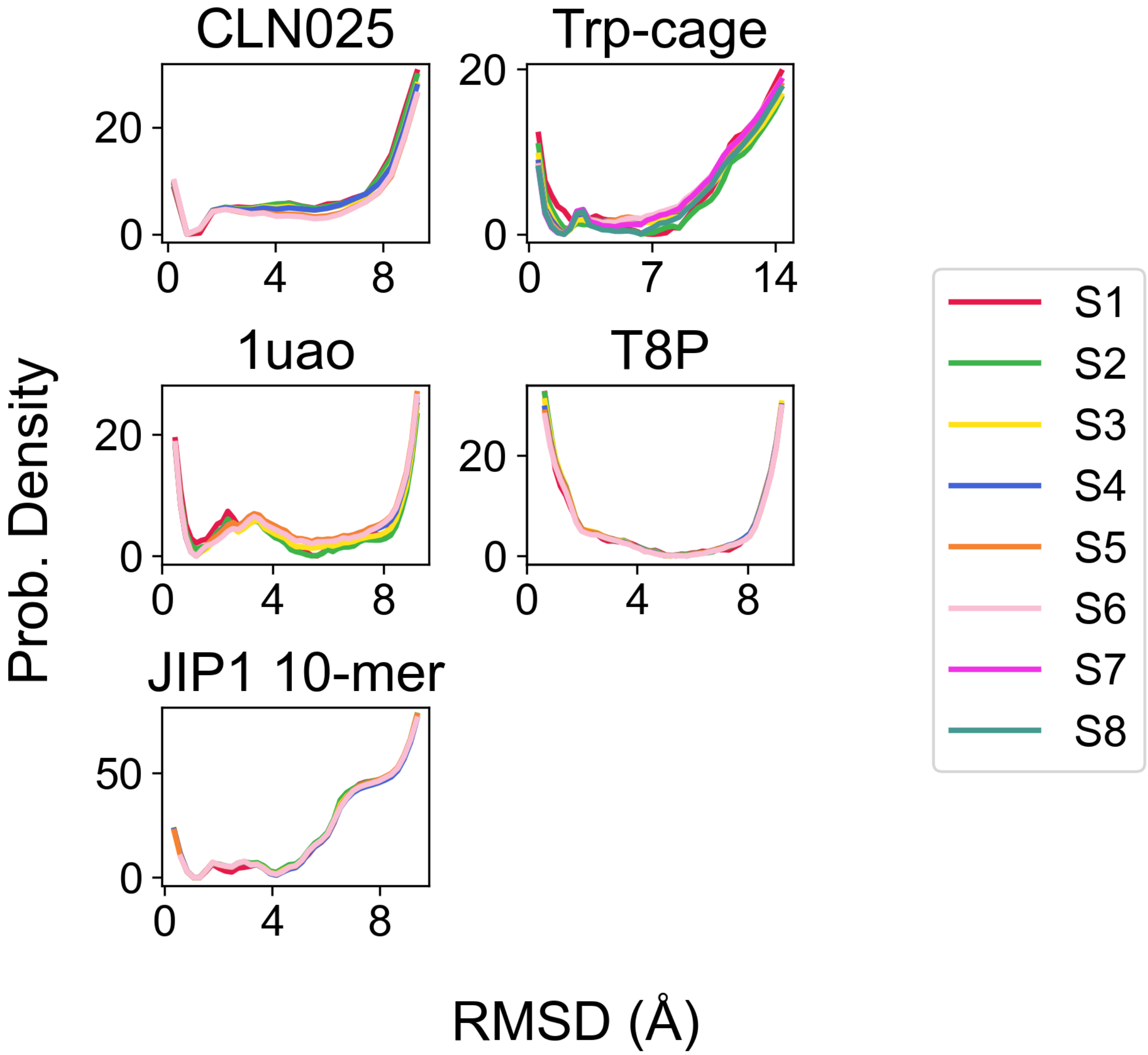


Figure S15: Converged free energy profiles for all proteins used to evaluate the SchNet implicit solvent model. Umbrella sampling windows were split 5 ways to show that the free energy profiles were converged. For CLN025, 1uao, and T8P, splits were created according the following scheme: S1 0 - 2 ns, S2 0 - 4 ns, S3 0 - 6 ns, S4 0 - 8 ns, S5 0 - 10 ns, and S6 0 - 13 ns. For the JIP1 10-mer, splits were created according to the following scheme: S1 0 - 5 ns, S2 0 - 10 ns, S3 0 - 15 ns, S4 0 - 20 ns, S5 0 - 25 ns, and S6 0 - 30 ns. For Trp-cage, splits were created according to the following scheme: S1 0 - 2.5 ns, S2 0 - 5 ns, S3 0 - 7.5 ns, S4 0 - 10 ns, S5 0 - 12.5 ns, S6 0 - 15 ns, S7 0 - 17.5 ns, and S8 0 - 20 ns. For all proteins, the final split includes all configurations for each window.

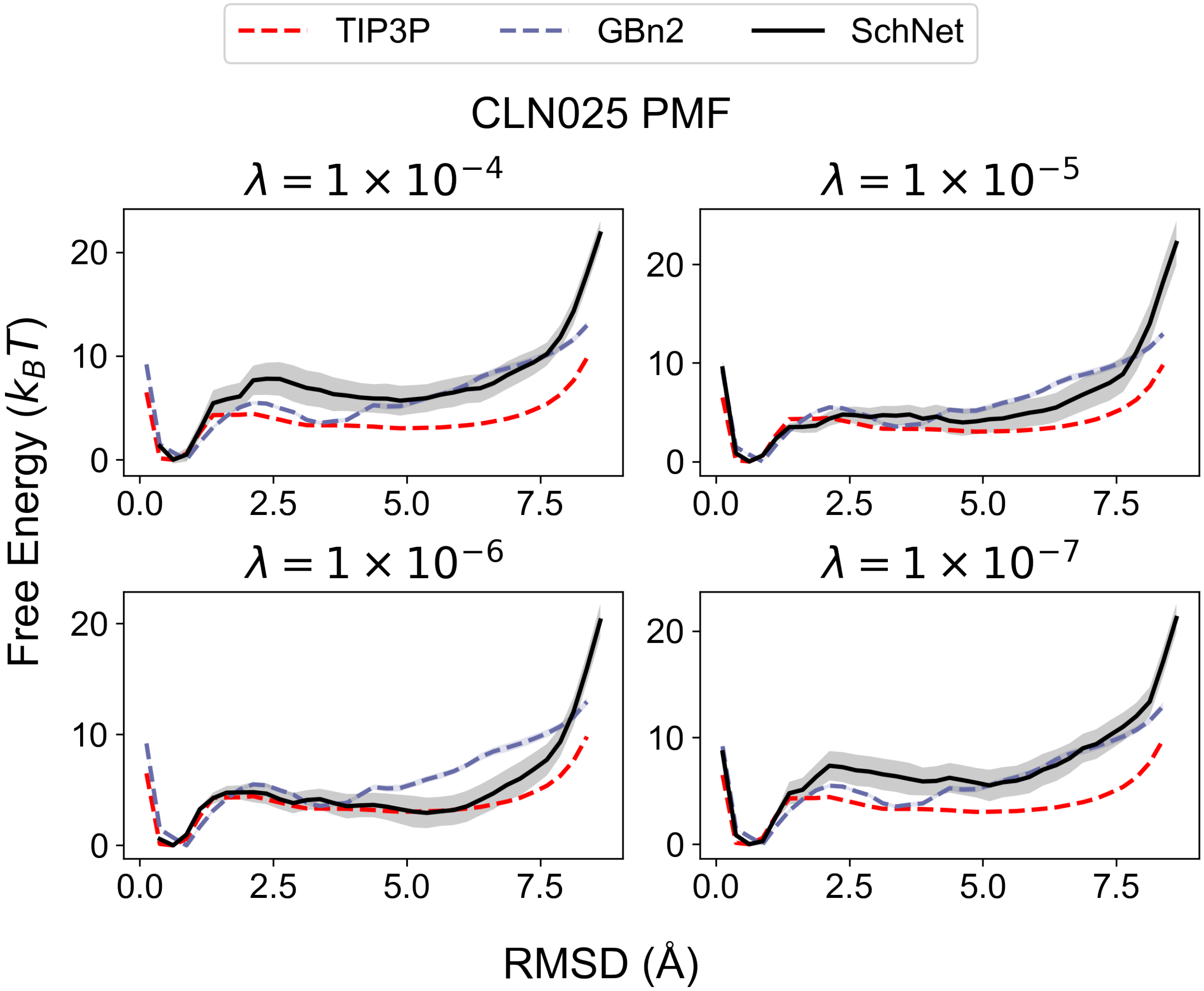


Figure S16: Impact of the L2 regularization constants, $\lambda$, on the performance of SchNet models ($N_{\text{IB}} = 3$, $r_{\text{cut}} = 1.8$ nm) trained with potential contrasting. We used the protein CLN025 as an example. Parameters in the interaction blocks were frozen during optimization with potential contrasting. The explicit solvent (TIP3P) free energy profile was computed from the configurations generated by Lindorff-Larsen et al.[S2], while configurations from umbrella sampling simulations were used to compute the GBn2 and SchNet free energy profiles. The SchNet model trained with $\lambda = 1 \times 10^{-6}$ produces a free energy profile that closely matches the explicit solvent result. See text *Section: Hyperparameters* for further discussion.

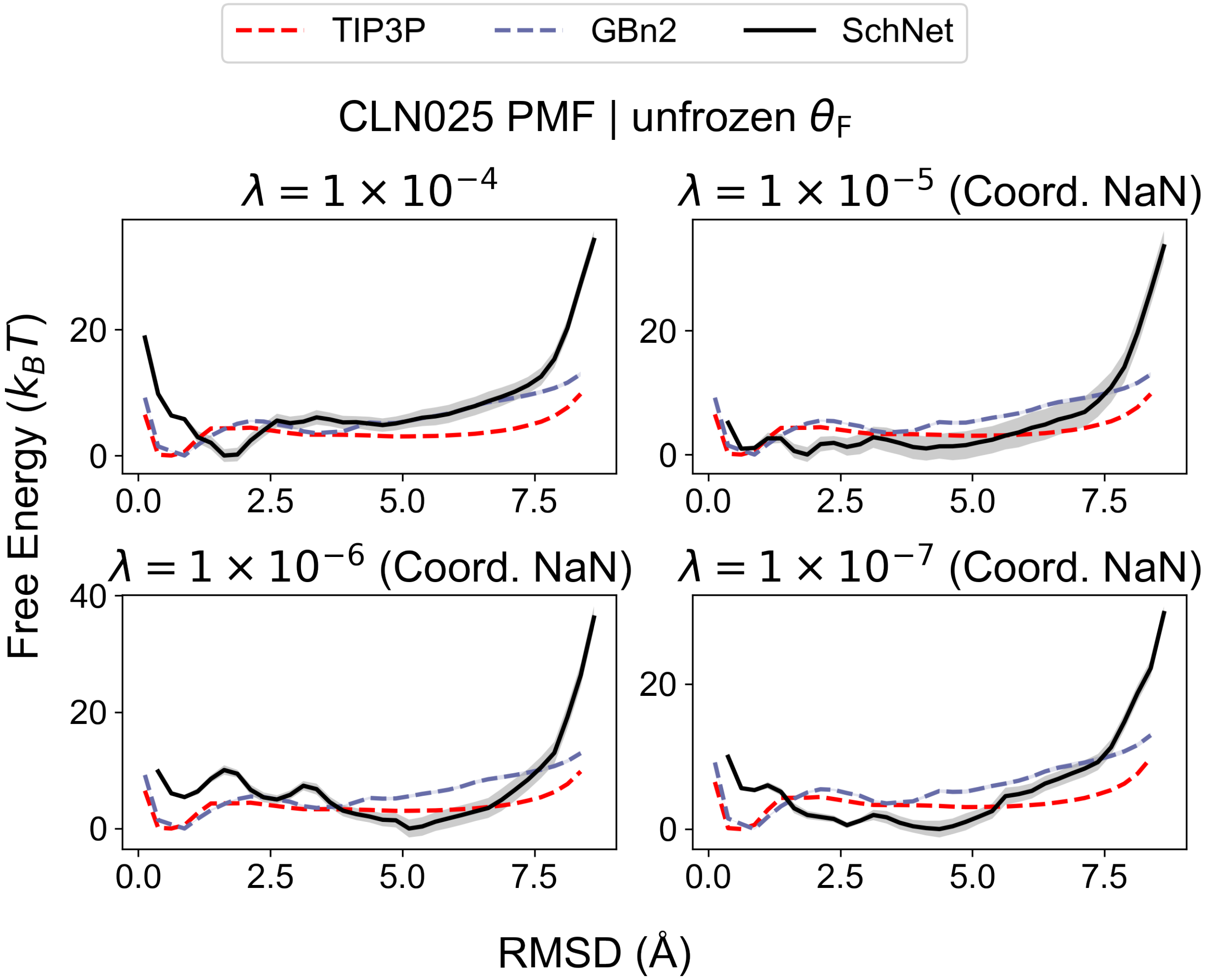


Figure S17: SchNet models ($N_{\text{IB}} = 3$, $r_{\text{cut}} = 1.8$ nm) trained using potential contrasting without freezing interaction block parameters performed poorly. We used the protein CLN025 as an example. The explicit solvent (TIP3P) free energy profiles was computed from the configurations generated by Lindorff-Larsen et al.[S2], while configurations from umbrella sampling simulations were used to compute the GBn2 and SchNet free energy profiles. No $\lambda$ resulted in a SchNet free energy profile that closely matched the explicit solvent result. SchNet models with $\lambda < 1 \times 10^{-4}$ result in simulation instability.

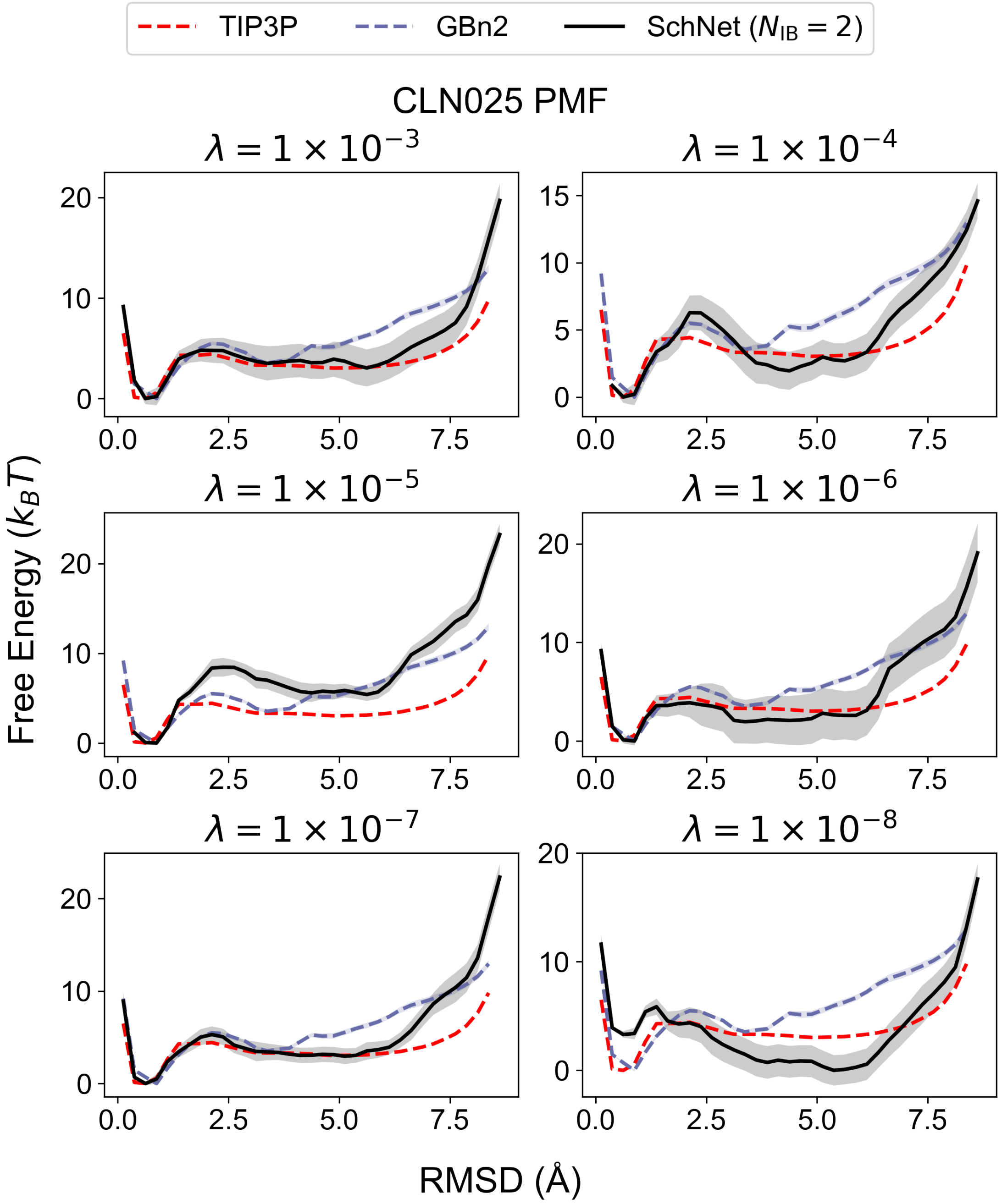


Figure S18: Impact of the L2 regularization constants, $\lambda$, on the performance of SchNet models ($N_{\rm IB} = 2$, $r_{\rm cut} = 1.8$ nm) trained with potential contrasting. We used the protein CLN025 as an example. The explicit solvent (TIP3P) free energy profiles was computed from the configurations generated by Lindorff-Larsen et al.[S2], while configurations from umbrella sampling simulations were used to compute the GBn2 and SchNet free energy profiles. The SchNet model with $\lambda = 1 \times 10^{-3}$ results in a free energy profile that most closely matches the explicit solvent result.

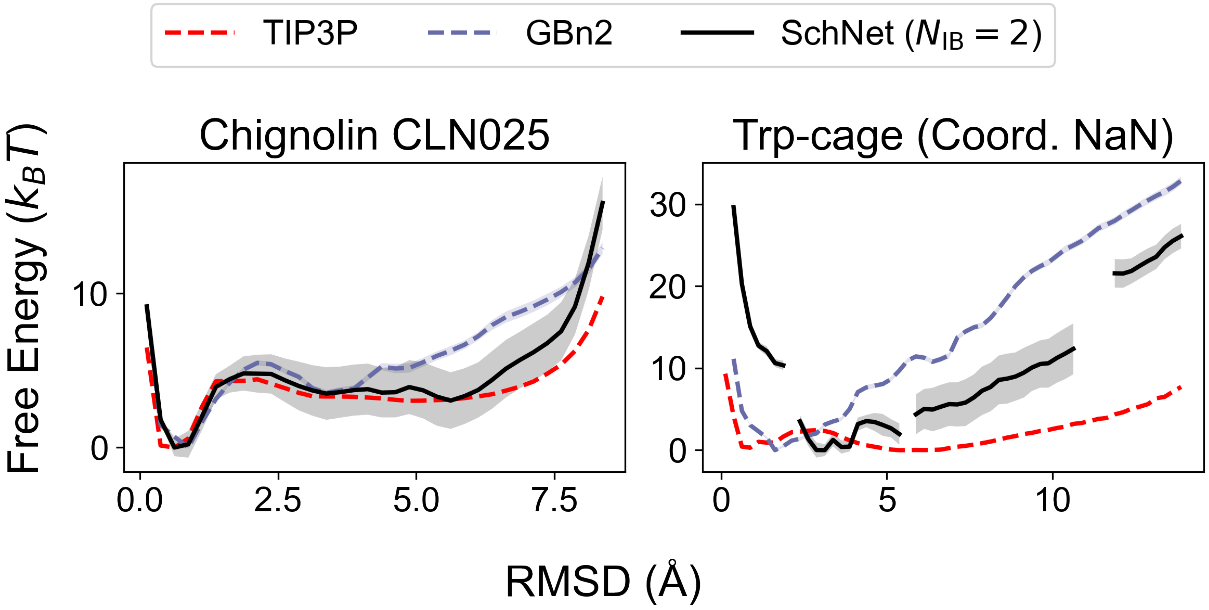


Figure S19: Free energy profiles for CLN025 and Trp-cage using a SchNet implicit solvent model with a smaller number of interaction blocks ($N_{\text{IB}} = 2$, $r_{\text{cut}} = 1.8$ nm, $\lambda = 1 \times 10^{-3}$). All explicit solvent (denoted as TIP3P) free energy profiles were computed from the configurations generated by Lindorff-Larsen et al.[S2], while configurations from umbrella sampling simulations were used to compute the GBn2 and SchNet free energy profiles. The SchNet free energy profile for CLN025 more closely matches the TIP3P free energy profile than the GBn2 free energy profile. Simulations for Trp-cage were too unstable to continue beyond 4 ns of production MD, and as such the Trp-cage SchNet free energy profile is not likely to be converged.

(S19) Paszke, A.; Gross, S.; Massa, F.; Lerer, A.; Bradbury, J.; Chanan, G.; Killeen, T.; Lin, Z.; Gimelshein, N.; Antiga, L.; Desmaison, A.; Kopf, A.; Yang, E.; DeVito, Z.; Raison, M.; Tejani, A.; Chilamkurthy, S.; Steiner, B.; Fang, L.; Bai, J.; Chintala, S. PyTorch: An Imperative Style, High-Performance Deep Learning Library. Advances in Neural Information Processing Systems. 2019.

(S20) Fey, M.; Lenssen, J. E. Fast Graph Representation Learning with PyTorch Geometric. *ArXiv* **2019**, preprint arXiv:1903.02428.

(S21) McGibbon, R. T.; Beauchamp, K. A.; Harrigan, M. P.; Klein, C.; Swails, J. M.; Hernández, C. X.; Schwantes, C. R.; Wang, L.-P.; Lane, T. J.; Pande, V. S. MDTraj: A Modern Open Library for the Analysis of Molecular Dynamics Trajectories. *Biophys. J.* **2015**, *109*, 1528–1532.

(S22) Schütt, K. T.; Sauceda, H. E.; Kindermans, P.-J.; Tkatchenko, A.; Müller, K.-R. SchNet – A deep learning architecture for molecules and materials. *J. Chem. Phys.* **2018**, *148*, 241722.

(S23) Kingma, D. P.; Ba, J. Adam: A Method for Stochastic Optimization. *ArXiv* **2017**, preprint arXiv:1412.6980.

(S24) Shirts, M. R.; Chodera, J. D. Statistically optimal analysis of samples from multiple equilibrium states. *J. Chem. Phys.* **2008**, *129*, 124105.

(S25) Tan, Z.; Gallicchio, E.; Lapelosa, M.; Levy, R. M. Theory of binless multi-state free energy estimation with applications to protein-ligand binding. *J. Chem. Phys.* **2012**, *136*, 144102.

(S26) Ding, X.; Vilseck, J. Z.; Brooks, C. L. A Fast Solver for Large Scale Multistate Bennett Acceptance Ratio Equations. *J. Chem. Theory Comput.* **2019**, *15*, 799–802.

(S27) Kunsch, H. R. The Jackknife and the Bootstrap for General Stationary Observations. *Ann. Stat.* **1989**, *17*, 1217–1241.

(S28) Virtanen, P.; Gommers, R.; Oliphant, T. E.; Haberland, M.; Reddy, T.; Cournapeau, D.; Burovski, E.; Peterson, P.; Weckesser, W.; Bright, J.; van der Walt, S. J.; Brett, M.; Wilson, J.; Millman, K. J.; Mayorov, N.; Nelson, A. R. J.; Jones, E.; Kern, R.; Larson, E.; Carey, C. J.; Polat, İ.; Feng, Y.; Moore, E. W.; VanderPlas, J.; Laxalde, D.; Perktold, J.; Cimrman, R.; Henriksen, I.; Quintero, E. A.; Harris, C. R.; Archibald, A. M.; Ribeiro, A. H.; Pedregosa, F.; van Mulbregt, P. SciPy 1.0: fundamental algorithms for scientific computing in Python. *Nat Methods* **2020**, *17*, 261–272.

(S29) Scott, D. W. *Multivariate density estimation: theory, practice, and visualization*; John Wiley & Sons, 2015.
